# Mapping gene flow between ancient hominins through demography-aware inference of the ancestral recombination graph

**DOI:** 10.1101/687368

**Authors:** Melissa J. Hubisz, Amy L. Williams, Adam Siepel

## Abstract

The sequencing of Neanderthal and Denisovan genomes has yielded many new insights about interbreeding events between extinct hominins and the ancestors of modern humans. While much attention has been paid to the relatively recent gene flow from Neanderthals and Denisovans into modern humans, other instances of introgression leave more subtle genomic evidence and have received less attention. Here, we present an extended version of the ARGweaver algorithm, ARGweaver-D, which can infer local genetic relationships under a user-defined demographic model that includes population splits and migration events. This Bayesian algorithm probabilistically samples ancestral recombination graphs (ARGs) that specify not only tree topology and branch lengths along the genome, but also indicate migrant lineages. The sampled ARGs can therefore be parsed to produce probabilities of introgression along the genome. We show that this method is well powered to detect the archaic migration into modern humans, even with only a few samples. We then show that the method can also detect introgressed regions stemming from older migration events, or from unsampled populations. We apply it to human, Neanderthal, and Denisovan genomes, looking for signatures of older proposed migration events, including ancient humans into Neanderthal, and unknown archaic hominins into Denisovans. We identify 3% of the Neanderthal genome that is putatively introgressed from ancient humans, and estimate that the gene flow occurred between 200-300kya. We find no convincing evidence that negative selection acted against these regions. We also identify 1% of the Denisovan genome which was likely introgressed from an unsequenced hominin ancestor, and note that 15% of these regions have been passed on to modern humans through subsequent gene flow.

## Introduction

It is well established that gene flow occurred among various ancient hominin species over the past several hundred thousand years. The most well-studied example is the interbreeding that occurred when humans migrated out of Africa and came into contact with Neanderthals in Eurasia roughly 50,000 years ago [1, 2]. This left a genetic legacy in modern humans which persists today: between 1-3% of the DNA of non-African humans can be traced to Neanderthals [3]. We also now know that an extinct sister group to the Neanderthals, the Denisovans, intermixed with humans in Asia, leaving behind genomic fragments in 2-4% of the DNA of modern Oceanian humans [4, 5].

Many other admixture events have been hypothesized, creating a complex web of ancient hominin interactions across time and space. These include: between Neanderthals and Denisovans (Nea↔Den) [2, 6]; between Neanderthals and ancient humans who left Africa over 100 thousand years (Hum→Nea) [7]; between an unknown diverged or “super-archaic” hominin (possibly *Homo erectus*) and Denisovans (Sup→Den) [2, 8]; and between other unknown archaic hominins and various human populations in Africa (Sup→Afr) [9, 10]. (In the above notation, the arrows indicate the direction of gene flow hypothesized; in many cases it may have gone both ways, but we lack samples to test the other direction).

As the network of interactions gets more complex, it becomes more difficult to apply standard methods to test for gene flow or identify introgressed regions [11]. For example, a positive value has been observed for the statistic *D*(*Neaderthal, Denisovan, African, Chimp*) [2], indicating that there is excess allele sharing between Neanderthals and African humans, as compared to Denisovans and Africans. But, this could be explained by gene flow between Neanderthals and Africans, or from super-archaic hominins into Denisovans, or some combination. The main strategy for teasing apart these scenarios is to examine the age of shared alleles. In this case, the *D* statistic is highest at sites where the derived allele is fixed or high-frequency in Africa, implying that many of the excess shared alleles are older than the Neanderthal/human divergence, so cannot be explained by Hum↔Nea gene flow. This forms the basis for the hypothesis of super-archaic introgression into Denisovans [2], which predicts a deficit of African-Denisovan shared alleles, as opposed to a surplus of African-Neanderthal sharing. However, it has also been noted that many genomic windows with the lowest Neanderthal-Africa divergence nevertheless have high Neanderthal-Denisovan divergence, which is best explained by Hum→Nea gene flow [7]. Currently, both events have support from multiple studies, including: model-based demography estimation by GPhoCS [7, 12], using ARGweaver [13] to examine coalescence times for gene trees that do not match the species tree [7], and comparing the frequency-stratified D-statistics with those from extensive simulations under various models of gene flow [8].

While both Sup→Den and Hum→Nea events have substantial support, it remains challenging to identify introgressed genomic regions that result from them. This problem is more difficult than identifying regions introgressed into modern non-African humans from Neanderthals and Denisovans, both because we do not have a sequence from the super-archaic hominin, and because these events are likely older, and therefore the haplotypes more broken up by recombination. We are further limited by the very small numbers of sequenced Neanderthal and Denisovan genomes. Current approaches, including the conditional random field (CRF) [3, 5] and the S* statistic [14, 15] (and recent variant Sprime [16]), have been tuned to the problem of finding recent introgression into humans. Furthermore, they only use a small number of summary statistics, such as locations of specific patterns of allele sharing. When the genomic signal is more subtle, it may be necessary to incorporate all the data with careful methodology in order to have sufficient power to confidently detect these regions.

In this paper, we present ARGweaver-D, which infers ancestral recombination graphs (ARGs) [17–19] conditional on a generic demographic model that includes population splits, size changes, and migration events. The ARG consists of local trees across a chromosome, representing the ancestral relationships among a set of sequenced individuals at every genomic position. In this extension to ARGweaver, the ARGs also contain information about the population membership of each lineage at every time point, so that introgressed regions are encoded in the ARG as lineages that follow a migrant path. Unlike most other methods, this approach allows multiple types of introgression to be inferred simultaneously, and takes into account the full haplotype structure of the input seqeunces. It works on unphased genomes and can accommodate changing migration and recombination rates. ARGweaver-D is a Bayesian method, using Markov chain Monte Carlo (MCMC) iterations to remove and “rethread” branches into the local trees; as a result, the output of ARGweaver-D is a series of ARGs that are sampled from the posterior distribution of ARGs conditional on the input data and demographic model. From these, we can extract posterior probabilities of introgression for any lineage at any genomic position.

Another recent method, dical-admix [20], is similar to ARGweaver-D in that it is designed to accommodate generic demographic models, and takes the full haplotype structure of the input sequences into account. However, there are some important differences that make our approach more applicable to the complex history of ancient hominins. dical-admix assumes that there are only a few admixed individuals, and that other genomes are “trunk” lineages that help define the haplotype structure of their respective populations. It therefore cannot infer admixture from an unsampled population, nor is it designed to work when all individuals have some degree of admixed ancestry. Additionally, ARGweaver-D can handle unphased genomes, which is important since there are not enough Neanderthal or Denisovan samples to reliably phase these archaic genomes.

After introducing ARGweaver-D, we present simulation studies showing it can successfully detect Nea→Hum introgression, even when using a limited number of genomes. We then use simulations to show that it can also detect older migration events, including Hum→Nea, Sup→Den, and Sup→Afr, depending on the underlying demographic parameters. We then apply this method to African humans and ancient hominins, classifying 3% of the Neanderthal genome as introgressed from ancient humans, and 1% of the Denisovan genome as introgressed from a super-archaic hominin. In contrast to Nea→Hum introgression, we do not see any evidence of selection against Hum→Nea introgression.

## Results

### ARGweaver-D can estimate genealogies conditional on arbitrary demographic model

ARGweaver-D is an extension of ARGweaver [13]that can infer ARGs conditional on a user-defined population model. This model can consist of an arbitrary number of present-day populations that share ancestry in the past, coalescing to a single panmictic population by the most ancestral discrete time point. Population sizes can be specified separately for each time interval in each population. Migration events between populations can also be added; they are assumed to occur instantaneously, with the time and probability defined by the user.

Recall that ARGweaver is a MCMC sampler, in which each iteration consists of removing a branch from every local tree in the ARG (“unthreading”), followed by the “threading” step, which resamples the coalescence points for the removed branches. This threading step is the main engine behind ARGweaver, and is accomplished with a hidden Markov model (HMM), in which the set of states at a particular site consists of all possible coalescence points in the local tree. In the original version of ARGweaver (with a single panmictic population), each of these states is defined by a branch and time. In ARGweaver-D, each state has a third property, which we call the “population path”, representing the population(s) assigned to the new branch throughout its time span. The modified threading algorithm is illustrated and further described in Fig 1.

**Fig 1.**
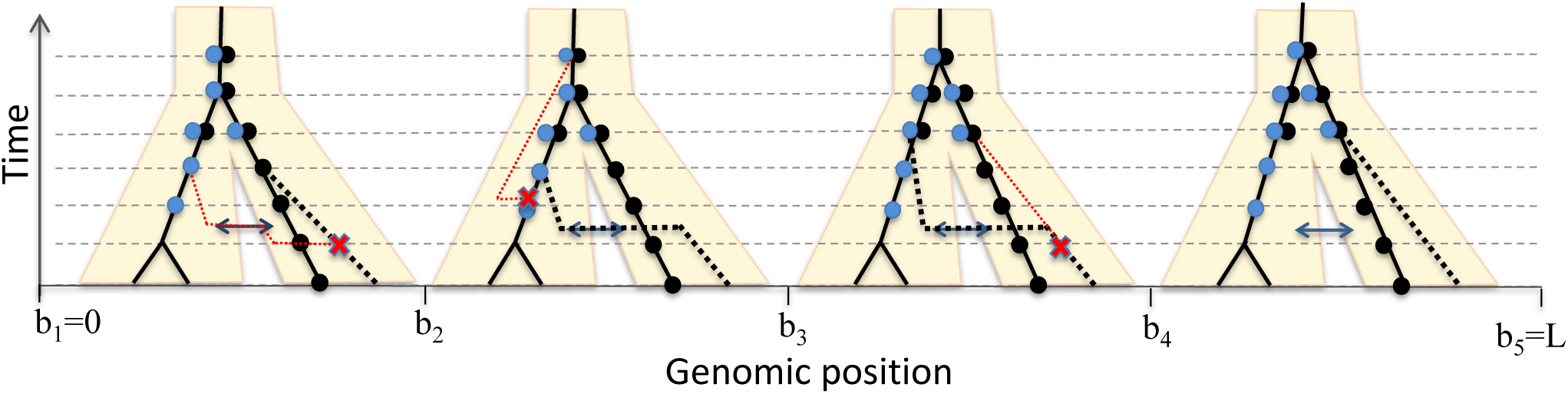
Illustration of the “threading” operation under a model with two populations and a single migration band. The gray horizontal dashed lines represent the discrete time points in the ARGweaver model, when coalescence and recombination events occur; migration and population divergence times are pre-specified by the user and rounded to the nearest “half time-point” (midway between the dashed lines). Migration is assumed to occur instantaneously at a rate *p_M_* specified by the uesr. This ARG currently has three haploid samples, indicated by the solid black lines. A fourth sample from the right-hand population is being threaded into the ARG, with the dotted black line representing one possible threading outcome. Each dot on the tree is a potential coalescence point for the new branch, representing a state in the threading HMM. The black dots are states from a population path with no migration, whereas the blue dots are from the migrant population path. is more efficient than the original panmictic Recombination events occur immediately before positions *b*_2_, *b*_3_, and *b*_4_, as indicated by the red X on the trees preceding those positions. The dotted red line shows the recoalescence of the broken branch, which defines the tree at the next site. The recombinations before *b*_2_ and *b*_4_ would be sampled after the threading algorithm, as they occur on the branch being threaded, whereas the recombination before *b*_3_ is part of the ARG before the threading, and therefore not modified at this stage. Here, we only show a single tree in each interval between recombination events; the local tree is identical within each of these intervals. The lineage being threaded enters an introgressed state at position *b*_2_, and leaves it at *b*_4_. The transition probabilities of the HMM are calculated between each pair of adjacent states; the probability of migration *p_M_* is a factor in the transition observed at *b*_2_. It is not a factor at *b*_3_ because the new branch is already in a migrant state. The transition probability at *b*_4_ includes a factor of 1 *− pM*.

Without migration events, and assuming that present-day population assignment for each branch is known, the ARGweaver-D version. This is because coalescence is not possible unless two branches are in the same population at the same time, and so the state space of potential coalescence points will be a subset of the original state space. However, as migration events are added, coalescence points in other populations become possible, and some coalescence points may be reachable by multiple population paths (see Fig 1). Therefore, the complexity of the algorithm can quickly increase. Whereas the original threading algorithm had an asymptotic running time of *O*(*Lnk*^2^) (where *L* is the number of sites, *n* the number of samples, and *k* the number of time points), ARGweaver-D is *O*(*Lnk*^2^*P* ^2^), where *P* is the maximum number of population paths available to any single lineage.

One way to improve the efficiency is to allow at most one migration event at any genomic location. Note that this assumption still allows multiple lineages to be introgressed at the same genomic position, if they are descended from a common migrant ancestor. This assumption is reasonable when the number of samples is small and the migration rate is low, and is set as a default in ARGweaver-D that we use throughout this paper. It has two advantageous side-effects: it avoids strange parts of the state space that could cause MCMC mixing problems (such as back-migrations, or population label switching issues). It also means that if we are modelling introgression from a ”ghost” population such as a super-archaic hominin (from which we have no samples), there will be at most one (migrant) lineage in the population at any location. Therefore, the population size of ghost populations does not matter as coalescence will not occur within them.

After running ARGweaver-D, it is straightforward to identify predicted introgressed regions; they are encoded in each ARG as lineages that follow a migration band. By examining the set of ARGs produced by the MCMC sampler, ARGweaver-D can compute posterior probabilities of introgression across the genome; this can be done in any way that the user would like: as overall probabilities of migration anywhere in the tree, or probabilities of a specific sampled genome having an ancestral lineage that passes through a particular migration band. For a diploid individual, we can look at probabilities of being heterozygously or homozygously introgressed. Throughout this paper we use the cutoff of *p* ≥ 0.5 to define predicted introgressed regions, and compute total rates of called introgression for a diploid individual as the average amount called across each haploid lineage.

More details of the ARGweaver-D algorithm are given in Fig 1 and the Supplementary Text. ARGweaver-D is built into the ARGweaver source code, which is available at: http://github.com/CshlSiepelLab/argweaver.

### ARGweaver-D can accurately identify archaic introgression in modern humans

We performed a set of simulations to assess the performance of ARGweaver-D for identifying Neanderthal introgression into modern humans. These simulations realistically mimic human and archaic demography, as well as variation in mutation and recombination rates (see Methods). We compared the performance with the CRF algorithm [3]; Fig 2 summarizes the results. Overall, ARGweaver-D has improved performance over the CRF, which is subtle for long segments but becomes more pronounced for shorter segments. This gain in power is despite the fact that the CRF used a much larger panel of African samples than was used by ARGweaver-D. (CRF used 43 Africans, ARGweaver-D used only 2 to save computational cost).

**Fig 2.**
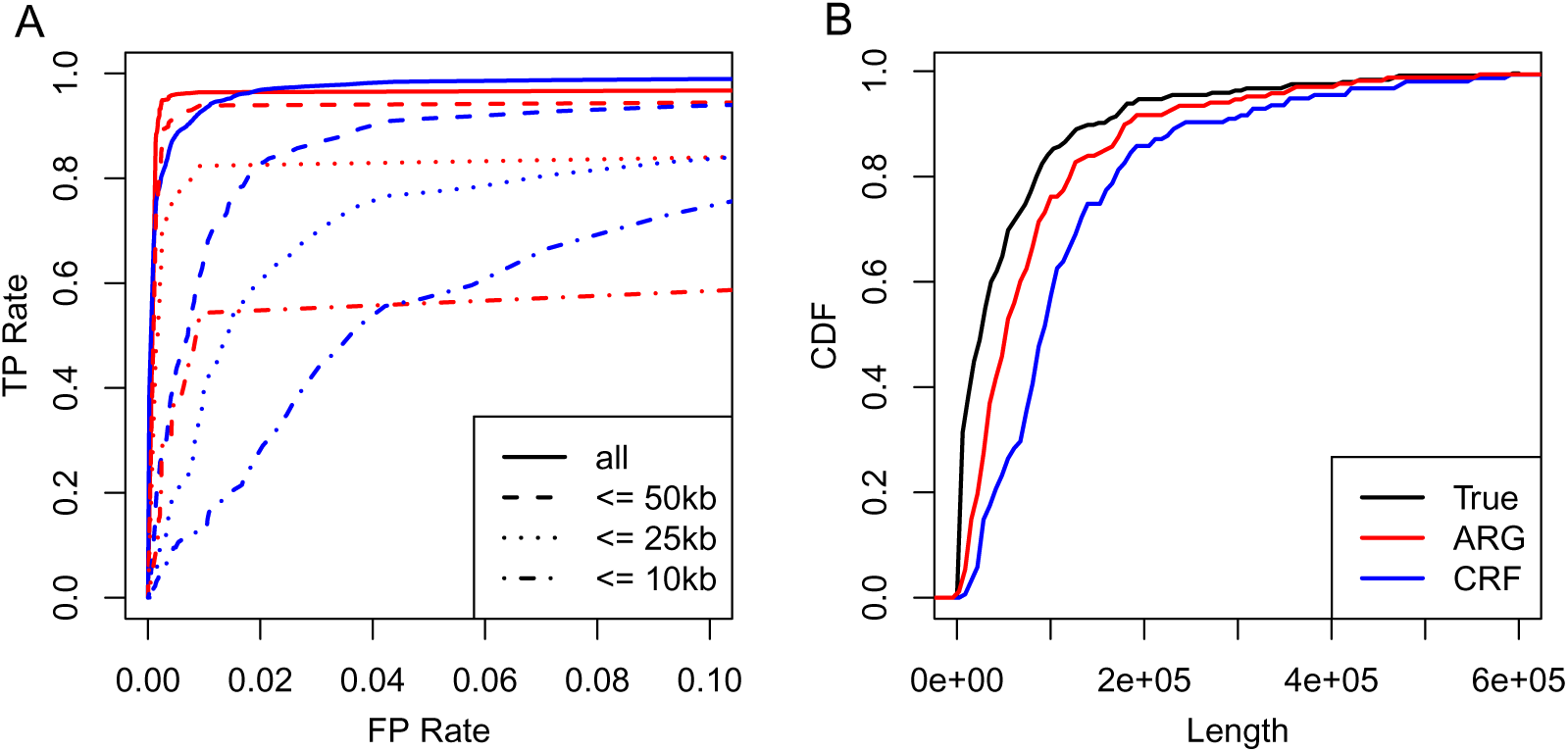
Performance on Nea→Hum simulations. **A:** ROC curves showing basewise performance of ARGweaver-D (red) and the CRF (blue) for predicting introgressed regions in simulated data. The two methods predicted introgression in the same simulated European individuals, however the CRF made use of the full reference panel (43 diploid Africans), whereas ARGweaver-D only used a small subset of the reference panel (2 diploid Africans). Different line patterns correspond to different maximum segment lengths. **B:** The length distribution of real and predicted introgressed regions for the same simulations and predictions shown in panel A.

Next, we predicted introgressed regions in two non-African human samples from the Simons Genome Diversity Panel (SGDP), one European (Basque) and a Papuan. The ARGweaver-D model used is illustrated in Fig 3; in this case only the “Recent migration” bands were included. We compared to calls on the same individuals from the CRF. Again, ARGweaver-D used two Africans, whereas the CRF used 43. And while the CRF uses Africans as a control group, ARGweaver-D allows for introgression into any of the human samples. The results are summarized in Fig 4. Overall, the two methods call a large fraction of overlapping regions, but each method also produces a substantial fraction not called by the other method (between 15-40%), and ARGweaver-D generally calls more regions. While ARGweaver-D seems to have greater power in simulations, another factor in the discrepancy may be that the ARGweaver-D segments were called with the inclusion of both the Altai and Vindija Neanderthals, whereas the CRF calls were produced with the Altai Neanderthal only. Both methods show a strong depletion of introgression on the X chromosome, especially in the Basque individual.

**Fig 3.**
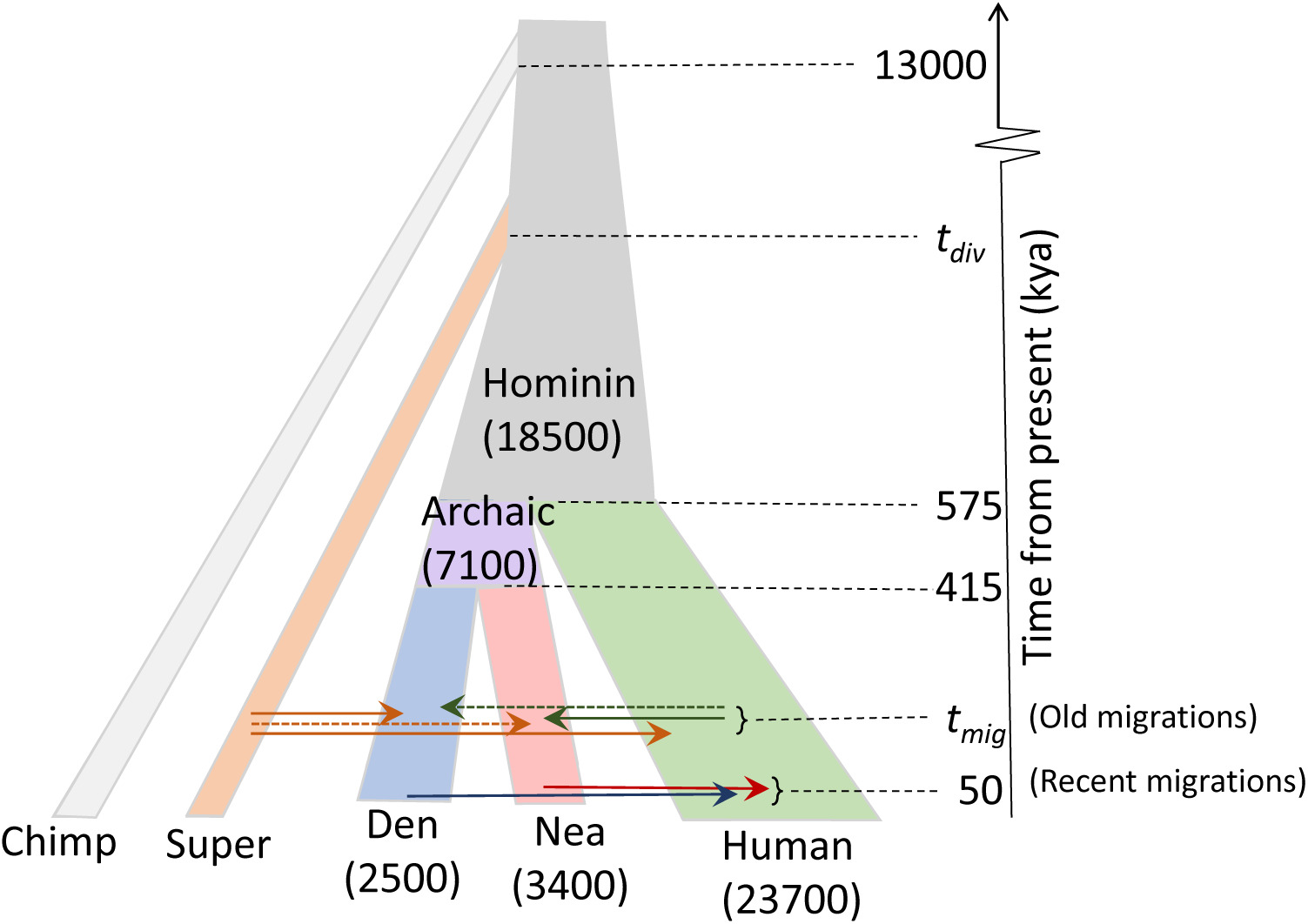
Population model used for ARGweaver-D analysis. Population sizes are given in parentheses. The model is invariant to the population sizes of the chimpanzee and super-archaic hominin, as no more than a single lineage ever exists in each of these populations. Migration events are shown by arrows between populations; solid arrows are used for events which have been proposed by previous studies. All parameters except *t_mig_* and *t_div_* are held constant at the values given.

**Fig 4.**
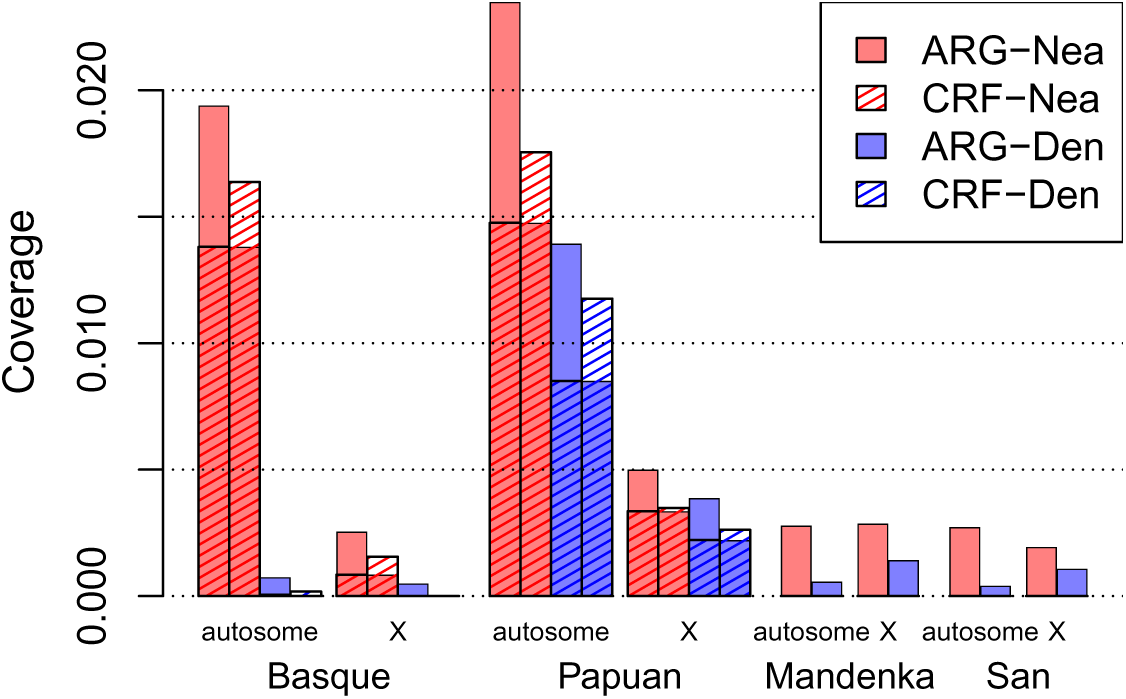
Average coverage of predicted introgressed regions into four SGDP individuals.s. The regions that are both colored and striped represent regions called by both CRF and ARGweaver-D. The CRF calls were only produced for non-African individuals, so only ARGweaver-D results are shown for Mandenka and San.

Notably, ARGweaver-D calls close to 0.5% introgression from Neanderthal into each of the African individuals. These calls may be explained by a combination of false positives and back-migration into Africa from Europe. Another possibility is that regions introgressed into Neanderthals from ancient humans [7] may be identified in the wrong direction under this model. With few samples, it is likely difficult to determine the direction of migration between two sister populations. Indeed, when we simulate migration in both directions, but still have only a Nea→Hum migration band in the ARGweaver-D model, 8% of Hum→Nea bases are identified as Nea→Hum. (See Supplementary Text). This is our motivation for excluding non-African samples when looking for introgressed regions from older migration events in the next section.

Finally, we compared the rate of calls in the Basque individual to predictions of Neanderthal ancestry based on the F4-Ratio statistic F4(Altai, chimp; Basque, African)/F4(Altai, chimp; Vindija, African) [11]. Both ARGweaver-D and CRF predicted fewer elements (1.95% and 1.56%, respectively), compared to the F4 ratio statistic (2.31%). Looking across the chromosomes, there is a higher correlation between coverage predicted by ARGweaver-D and the expectation from the F4 ratio (Spearman’s *ρ* = 0.75), than between CRF and the F4 ratio (*ρ* = 0.51) (Fig S1).

### ARGweaver-D can detect older introgression events

We next did a series of simulations to assess ARGweaver-D’s power to detect other ancient introgression events that have been previously proposed. To focus on these older events, we simulated the modern human samples using on a model of African human population history, and as such did not include the migration from Neanderthals or Denisovans into non-African humans. The simulations included three migration events: from modern humans into Neanderthals (Hum→Nea), from a “super-archaic” unsampled hominin into Denisovans (Sup→Den), and also from super-archaic into Africans (Sup→Afr). (Note that although both Sup→Afr and Sup→Den involve introgression from the same super-archaic population, it is only meant to represent introgression from any unsampled, diverged hominin species, and does not necessarily imply that the same population admixed with both Africans and Denisovans.) The simulations included many realistic features: ancient sampling dates for the archaic hominins, variation in mutation and recombination rates, randomized phase, and levels of missing data modeled after the SGDP and ancient genomes that we use for analysis (see Methods). Each set of simulations contained all three types of migration and ARGweaver-D detected all migration events in a single run with multiple migration bands in the model.

We analyzed these data sets with ARGweaver-D using the model depicted in Fig 3, with only the “old migration” bands. As we do not have good prior estimates for the migration time (*t_mig_*) or super-archaic divergence time (*t_div_*), we tried four values of *t_mig_* (50kya, 150kya, 250kya, 350kya) and two values of *t_div_* (1Mya, 1.5Mya). We generated data sets under all 8 combinations of *t_mig_* and *t_div_*, and then analyzed each data set with ARGweaver-D under all 8 models, in order to assess the effects of model misspecification on the inference.

The power of ARGweaver-D to detect introgression is summarized in Fig 5. The left side of the plot represents simulations generated with *t_div_* = 1Mya, whereas the right side used *t_div_* = 1.5Mya. Power to detect super-archaic introgression is clearly much higher when the divergence is higher, but (as expected) does not affect power to detect Hum→Nea introgression. Looking from top to bottom, the plots show the effect of increasing the true time of migration. In the top plot with *t_mig_* = 50kya, only results for Sup→Afr are shown because the archaic hominin fossil ages pre-date the migration time. For all events, we see power decrease as the true migration time decreases.

**Fig 5.**
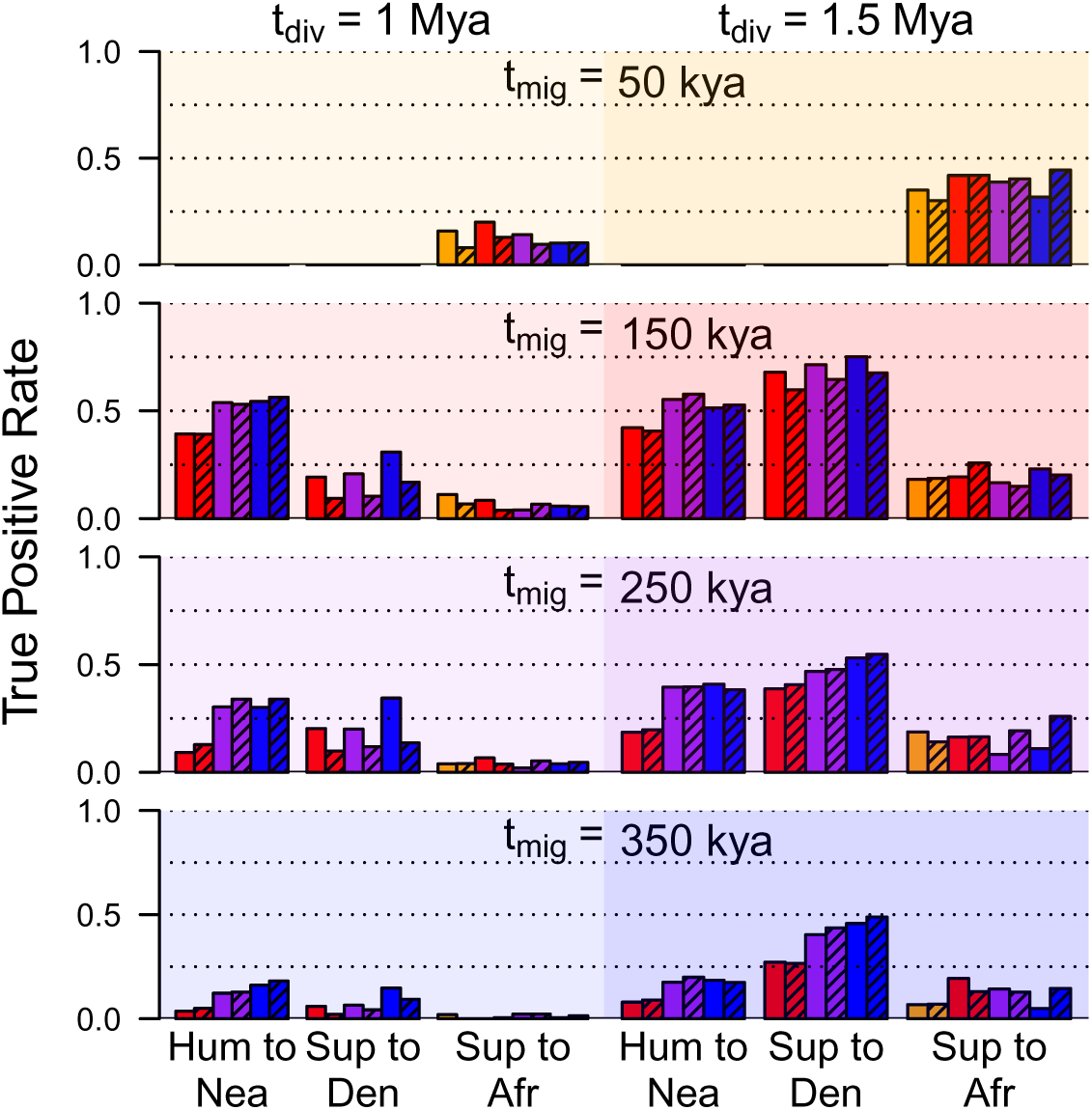
Simulation results. Each shaded box represents a set of simulations generated with a different value of *t_mig_* and *t_div_*. Each bar gives the basewise true positive rate for a particular migration event and ARGweaver-D model, using a posterior probability threshold of 0.5. The color of each bar represents the value of *t_mig_* used in the inference model (orange=50kya, red=150kya, purple=250kya, blue=350kya); shaded bars have *t_d_iv*=1.5Mya in the inference model, whereas solid bars use *t_div_* = 1.0Mya. Because the archaic hominin fossil ages are older than 50kya, results for *t_mig_* = 50kya (top) are only applicable for introgression into humans.

For a given simulation set, the effect of the parameters used by ARGweaver-D are generally more subtle. We note that power tends to be better when older migration times are used in the model, even when the true migration time is recent; in particular, the power when *t_mig_* = 150kya (red bars) is often much worse than when later times are used, especially for the Hum→Nea event. Similarly, power is often better when *t_div_* is set to 1Mya in the ARGweaver-D model, as opposed to 1.5Mya.

In summary, ARGweaver-D has reasonably good power to detect super-archaic introgression when the divergence time is old, but power is more limited as the divergence decreases. The power to detect Sup→Afr is always lower than the power to detect Sup→Den, as the African population size is much larger, making introgression more difficult to distinguish from incomplete lineage sorting. For the Hum→Nea event, we have around 50% power if the migration time is 150kya, and around 30% power when it is 250kya.

False positive rates are less than 1% when a posterior probability threshold of 0.5 is used (Fig S2). When analyzing the simulated data sets, we included two additional migration bands in the ARGweaver-D model as controls: one from the super-archaic population to Neanderthal (Sup→Nea), and another from humans into Denisova (Hum→Den). The rates of calling these events were also less than 1% for all models. Importantly, the rate of mis-classification is very low for all categories (Fig S3); in particular, the model can easily tell the difference between Hum→Nea and Sup→Den events, despite both resulting in similar *D* statistics [7, 8].

More details about the simulation results are available in the Supplementary Text. One issue to note is that, although the simulated data sets were generated with a human recombination map, the ARGweaver-D model used a simple constant recombination rate. Performance is somewhat better when ARGweaver-D uses the true recombination map, but in practice there are not enough Neanderthal or Denisovan samples to generate a reliable recombination map, and there is no data to infer the recombination map for the super-archaic population. The Supplementary Text also shows results when we simulate more African individuals. We find that performance does not improve as samples are added, so in the main text we focus on analysis with two African samples (four haploid genomes).

### Deep introgression results

We next applied the models from the previous section to real modern and archaic human genomes. Our goals were to identify and characterize introgressed regions from previously proposed migration events, as well as to see if we find evidence for other migrations which may not be detectable using other methods. Our data set consisted of two Africans from the SGDP [21], two Neanderthals [2, 8], the Denisovan [4], and a chimpanzee outgroup. We again use the demography illustrated in Fig 3, with old migration events only. We focus on the model with *t_mig_*=250kya and *t_div_*=1Mya, because this model seemed to have high power in all our simulation scenarios, and because our results suggest that it may be the most realistic (as discussed below). The results using other models are consistent with those presented here, and are described in the Supplementary Text.

An overview of the coverage of predicted introgressed regions is depicted in Fig 6; a more detailed summary is given in Fig S4. The most immediate observation is that Hum→Nea regions are called most frequently, at a rate of ∼3% in both the Altai and Vindija Neanderthal. This number is almost certainly an underestimate, given that the true positive rate for this model was measured between 30-55%. By contrast, only ∼0.37% of regions are classified as Hum→Den. As no previous study has found evidence for Hum→Den migration, this serves as a control, verifying that our false positive rate estimated in simulations is likely fairly accurate, as we estimated a FP rate of 0.41%.

**Fig 6.**
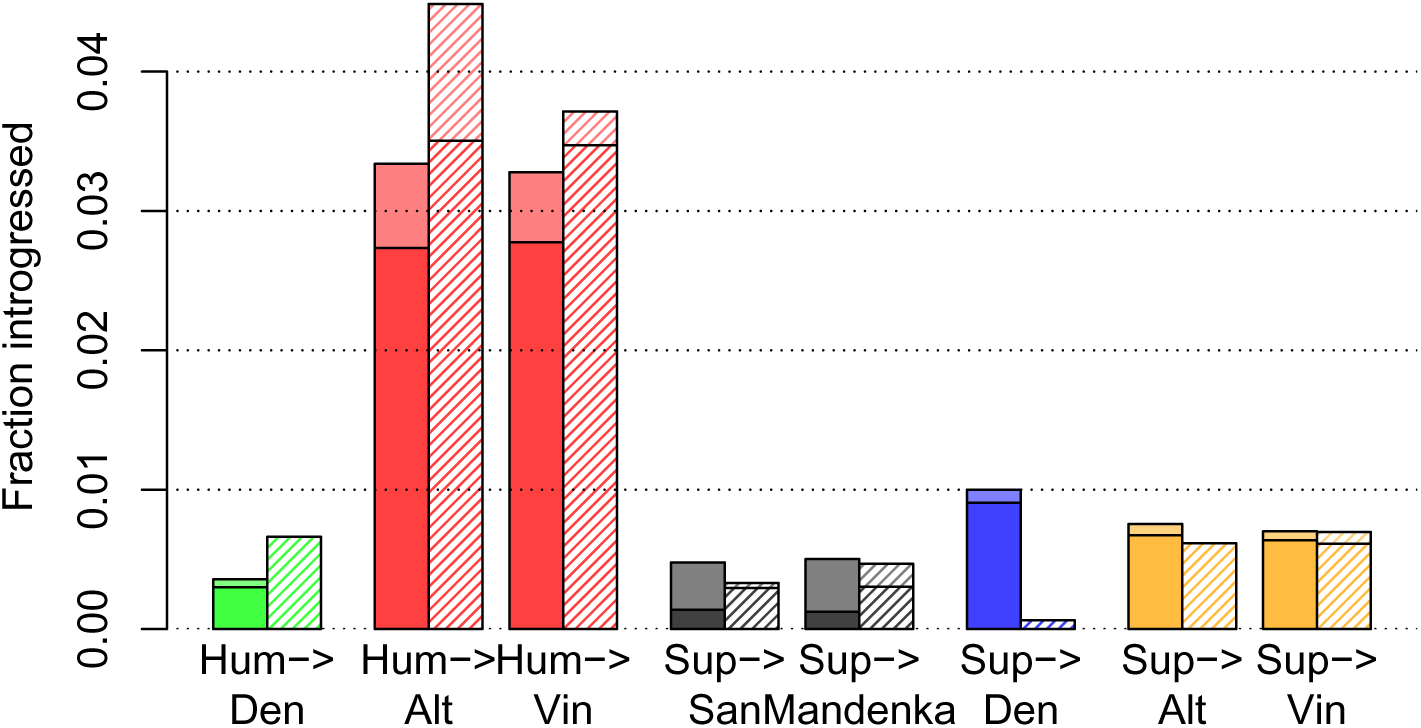
Genome-wide coverage of predicted ancient introgression. These results use a posterior probability cutoff of 0.5; solid bars are for autosomes, and striped bars for chromosome X. Each bar shows total average coverage for a haploid genome; the darker bottom portions of each bar represent homozygous calls.

Whereas there is a depletion on the X chromosome of archaic introgression into humans, we see high coverage of Hum→Nea on the X for both Altai and Vindija. The fact that it is somewhat higher on the X than the autosomes might be partially explained by increased power on the X; simulations suggest that power will be ∼20% higher for this event when population sizes are multiplied by 0.75 (Fig S5). Overall, there is a lot of variation in detected introgression across the chromosomes, and several autosomal chromosomes have higher predicted coverage than the X, including 1, 6, 21, and 22 (Fig S4).

Although the Vindija sample is 70ky younger than the Altai sample [8], there is no apparent depletion of human ancestry on Vindija compared to Altai on the autosomes, suggesting that negative selection did not cause a significant loss of human introgressed regions in the Neanderthal during that time. Several chromosomes do show drops in coverage from Altai to Neanderthal, with the largest drop on the X chromosome (Fig S4).

Other migrations are detected at lower levels. We identify 1% of the Denisovan genome as introgressed from a super-archaic hominin, which is double our estimated false positive rate for this event. The fact that we found much less than the ∼6% estimated by previous methods [8] might suggest that the super-archaic divergence time is closer to 1Mya, since we would expect to have more power with a higher divergence time. Still, this analysis resulted in 27Mb of sequence that may represent a partial genome sequence from a new archaic hominin. ARGweaver-D also predicted a small fraction of the Neanderthal genomes as introgressed from a super-archaic hominin (0.75% for Altai and 0.70% for Vindija). These amounts are only slightly above the estimated false positive rates (0.65%), and the Sup→Nea event has not been previously hypothesized.

One interesting aspect of Sup→Den and Sup→Nea regions is that, to the extent that these predictions are accurate, there is the potential that this super-archaic sequence was passed to modern humans through subsequent Den→Hum and Nea→Hum migrations. We explored these regions further by intersecting them with introgression predictions across the full SGDP data set. This analysis is detailed in the Supplementary Text. It first confirms that most Sup→Den and Sup→Nea regions have higher-than expected divergence to the Denisovans and Neanderthals (respectively) across all humans, and not just the two African humans used by ARGweaver-D. 15% of the Sup→Den regions overlap with sequence introgressed into Asian and Oceanian individuals from Denisovans, and many of these regions also contain a high number of variants consistent with super-archaic introgression. We also see that 35% of the Sup→Nea regions are introgressed in at least one modern-day non-African human. We also identified one region of hg19 (chr6:8450001-8563749) which appears to be Neanderthal-introgressed and overlaps a Sup→Nea region. We compiled a list of Sup→Den and Sup→Nea regions that overlap human introgressed regions, and the genes that fall in these regions. These are given in Tables S1 and S2.

**Table 1.** Amount of Hum→Nea introgression in deserts of Nea→Hum and Den→Hum introgression.

| Location (hg19) | cov Altai | cov Vindija | num Altai | num Vindija |
| --- | --- | --- | --- | --- |
| chr1:99-112 Mb | 0.097 | 0.024 | 4 | 3 |
| chr3:78-90 Mb | 0.101 | 0.078 | 6 | 5 |
| chr7:108-128 Mb | 0.023 | 0.031 | 4 | 5 |
| chr13:49-61 Mb | 0.029 | 0.048 | 6 | 5 |

The columns “cov Altai” and “cov Vindija” show the coverage of Hum→Nea within the given region on each Neanderthal; “num Altai” and “num Vindija” show the number of introgressed regions > 50kb. The genome-wide average coverage for Altai and Vindija is 0.034 and 0.033, respectively.

We examined lengths of all our sets of predicted regions, as they might be informative about the time of migration. However, we find that there is strong ascertainment bias towards finding longer regions, so that the length distributions are highly overlapping for different migration times. (See Supplementary Text).

Instead, we looked at the frequency spectrum of introgressed regions to gain insight into the times of migration events. The older the migration, the more likely that an introgressed region has drifted to high frequency and is shared across the sampled individuals. For the Hum→Nea event, we observed 37% of our regions are inferred as “doubly homozygous” (that is, introgressed across all four Neanderthal lineages). This is very close to what we observe in regions predicted from our simulations with migration at 250kya (38%), whereas simulations with migration at 150kya and 350kya had doubly-homozygous rates of 10% and 55%, respectively. To further narrow down the range of times, we did additional simulations with *t_mig_*=200, 225, 275, and 300kya, and compared the frequency spectrum of introgressed regions after ascertainment with ARGweaver-D. We find that the observed frequency spectrum is consistent with 200kya *< t_mig_ <* 300kya (Fig S6). The same approach suggests that *t_mig_ >* 225kya for the for the Sup→Den event (Fig S7).

### Data release and browser tracks

Our predictions and posterior probabilities can be viewed as a track hub on the UCSC Genome Browser [22], using the URL: http://compgen.cshl.edu/ARGweaver/introgressionHub/hub.txt. The raw results can be found in the sub-directory: http://compgen.cshl.edu/ARGweaver/introgressionHub/files. Fig 7 shows a large region of chromosome X as viewed on the browser, with a set of tracks showing called regions, and another showing posterior probabilities. Fig 8 shows a zoomed-in region with a Sup→Den prediction, and Fig S8 shows an example Hum→Nea region. When zoomed in, there is a track showing the patterns of variation in all the individuals used for analysis, with haplotype phasing sampled from ARGweaver-D.

**Fig 7.**
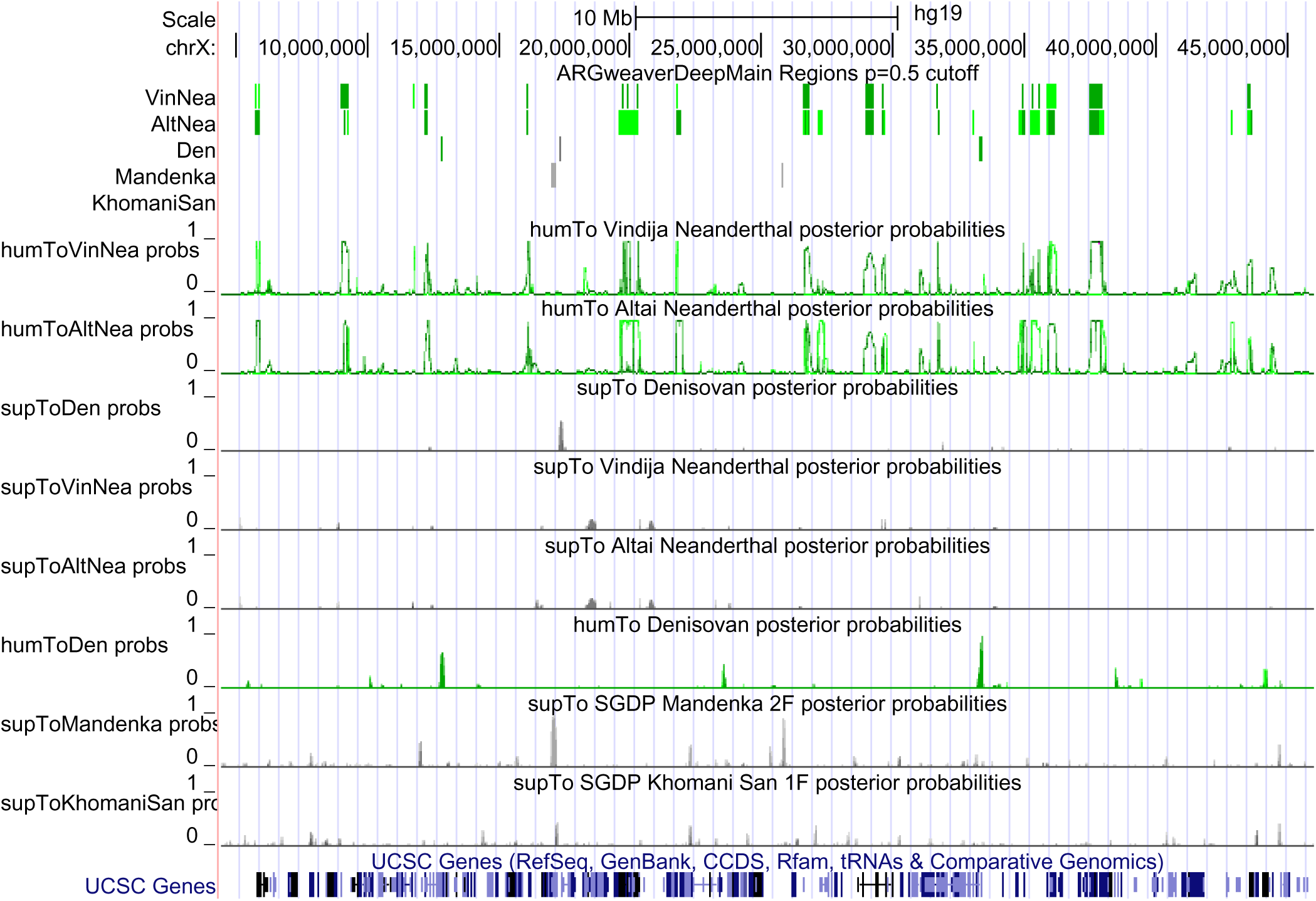
Introgression results for a large section of chromosome X displayed on UCSC Genome browser. The top set of tracks show predicted introgressed regions, with green indicating introgression from humans, and gray indicating introgression from a super-archaic hominin. Darker colors are used for homozygous introgression. Below that can be seen the posterior probabilities for each type of introgression into each individual.

**Fig 8.**
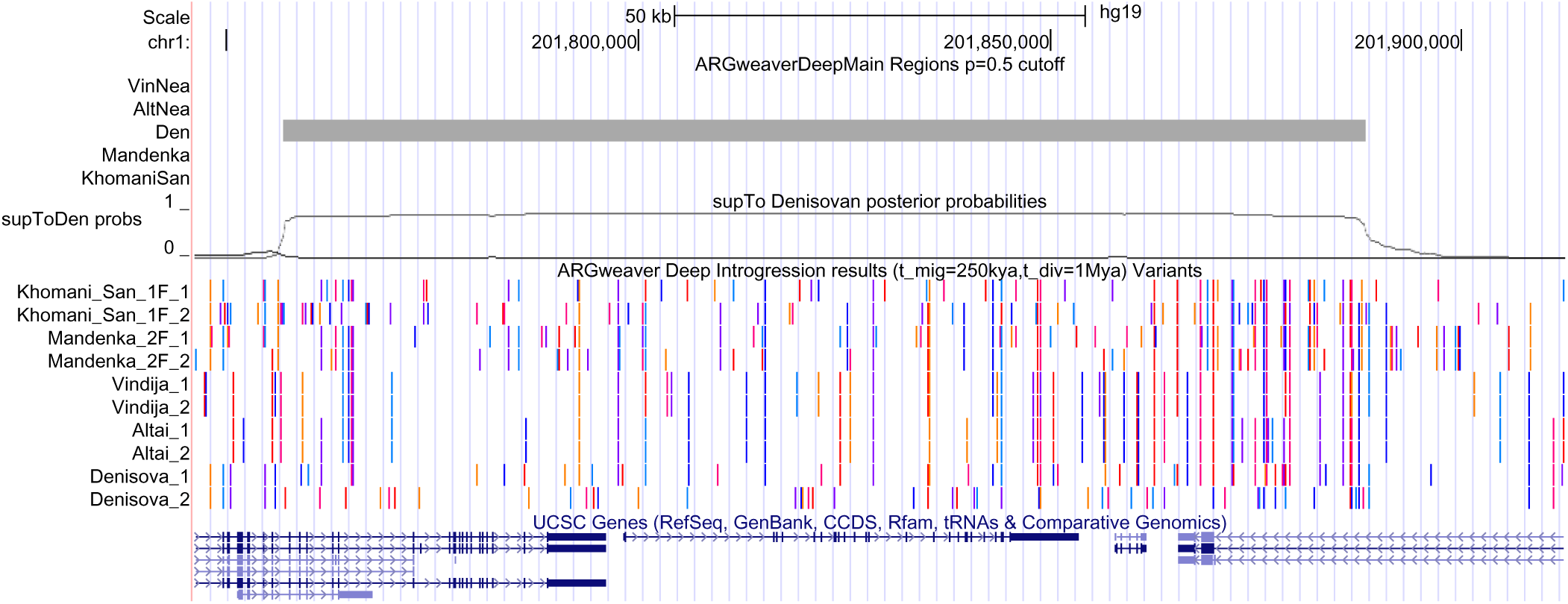
UCSC Genome Browser shot of a region with predicted heterozygous Sup→Den introgression. This browser shot is zoomed in on a 170kb region. The first two tracks show the predicted regions and posterior probabilties as in Fig 7, except only the supToDen probabilities are shown. The track just below shows the variants observed in this region that are used in the ARGweaver-D analysis. Alternating colors are used for each variant site. When chimpanzee alignments are available, the non-chimp allele is colored; otherwise the minor allele is colored. Lack of a color may mean that the haplotype has the chimpanzee or major allele, or that it has missing data. The phasing of the variants represents the final phase sampled by the ARGweaver-D algorithm. Here, the Denisovan is usually homozygous and shares variants with Africans and Neanderthals outside of the introgressed region; but within it, the Denisova 2 haplotype has many singleton variants, whereas Denisova 1 continues to share many variants with Neanderthals and Africans.

### Functional analysis of introgressed regions

Some observations in the previous section suggest that there was not strong selection against the Hum→Nea regions. We sought to look for other signals that might hint at possible functional consequences of this event.

We first looked at deserts of introgression that were detected in [5]. They noted four 10Mb deserts in which the rate of both Nea→Hum and Den→Hum introgression is *<* 1*/*1000. The coverage of Hum→Nea introgression within these deserts is shown in Table 1; the fairly high coverages suggest that these deserts are unidirectional. For two of the deserts, the Hum→Nea coverage is very high, especially in the Altai Neanderthal. The third region is interesting as it overlaps the FOXP2 gene, which contains two human-chimp substitutions that have been implicated in human speech [23, 24], although the Hum→Nea introgressed region is upstream of these substitutions (Fig S9).

We next looked at all deserts of Nea→Hum ancestry, to see if this larger set of regions are are depleted for introgression in the other direction. Based on the CRF regions, we identified 30 regions of at least 10Mb which qualify as deserts. We looked at several statistics, including coverage of Hum→Nea in these regions, number of elements, and change in coverage between the Altai and Vindija Neanderthals; but we do not see any difference in the distribution of these statistics within deserts, as compared to randomly chosen genomic regions matched for size (Fig S10).

Finally, we checked for enrichments or depletions of various functional elements in our introgressed segments, relative to what would be expected if the introgressed segments were randomly distributed throughout the genome. However, the interpretation of these numbers is difficult, as local genomic factors (such as effective population size, mutation and recombination rates) affect the power to detect regions. While the overall levels of enrichment are therefore difficult to interpret, it is interesting to note that the enrichment of functional regions (such as CDS, promoters, and UTRs) tends to be higher in the Altai than the Vindija Neanderthal, which is the opposite pattern we might expect from negative selection (since the Vindija Neanderthal’s fossil is much more recent). Further enrichment results are detailed in the Supplementary Text.

## Discussion

We present a new method for building ARGs under an arbitrary demographic model, and use this method as a powerful new way to identify introgressed regions. While it can detect introgressed Neanderthal and Densiovan sequences in human genomes, ARGweaver-D is too computationally complex to be used on a large scale over many human samples. However, it is very powerful even on small samples and older migration events, and has several other benefits over other methods. It does not require a reference panel of non-introgressed individuals, and can simultaneously identify introgression stemming from multiple migration events, as well as from both sampled or unsampled populations. ARGweaver-D does not rely on summary statistics, but uses a model of coalescence and recombination to generate local gene trees that are most consistent with the observed patterns of variation, even for unphased genomes. By incorporating all this information, it can successfully distinguish migration from incomplete lineage sorting, and tease apart different migration events that produce similar *D* statistics (such as Sup→Den and Hum→Nea). The code is freely available and can be applied to any number of species or demographic scenarios.

Applying this method to modern and archaic hominins, we confirm that a significant proportion of the Neanderthal genome consists of regions introgressed from ancient humans. While we identified 3% of the Neanderthal genome as introgressed, a rough extrapolation based on our estimated rates of true and false positives suggests that the true amount is around 6%. Thus, the Neanderthal genome was likely more influenced by introgression from ancient humans, than non-African human genomes are by Neanderthal introgression. Our follow-up analysis suggests that the Hum→Nea gene flow occurred between 200-300kya. This time estimate is largely based on the frequency of introgressed elements among the two diploid Neanderthal genomes, and thus will be sensitive to the accuracy of the demographic model we used for simulation, as well as other factors such as mutation rate and generation time.

Making conclusions about the possible impact of the Hum→Nea migration has proved challenging due to the myriad ascertainment biases—known and unknown—that affect our power to detect introgressed regions. Even in the case of Nea→Hum migration, in which power to detect introgression is much higher, earlier claims of depletion near genes, as well as decreasing levels of introgression over time, have been recently called into question [11, 20]. The strongest remaining pieces of evidence for negative selection against Nea→Hum introgression are the depletion on the X chromosome and several other genomic deserts. But for Hum→Nea, we see no depletion on the X, and while we do not have enough samples to detect deserts across Neanderthals, we confirm that previously identified Nea→Hum deserts are not depleted for introgression in the opposite direction. We do see a slight decrease in Hum→Nea introgression on the X chromosome in the Vindija Neanderthal compared to the Altai, which could be explained by weak negative selection removing some introgressed regions in the ∼70ky that separate these fossils. An interesting question is whether this lack of selection is because human introgression introduced healthy variation into the Neanderthal genome, or because the Neanderthal population was too small for natural selection to act against anything but the most harmful variants. However, without more archaic samples, these questions will be challenging to answer.

ARGweaver-D also identified 1% of the Denisovan genome as introgressed from a super-archaic hominin. Previous studies have estimated the total amount of Sup→Den as roughly 6% [8], but this is the first study to be able to identify specific introgressed regions. The fact that we only find a small fraction of the total amount suggests that the introgressing population was not too highly diverged from other hominins; this low power is much more consistent with a divergence time of 1Mya than 1.5Mya. Still, we report 27Mb of putative super-archaic sequence from this previously-unsequenced hominin, and we note that 15% of these regions have been passed on to modern humans through Den→Hum introgression. It may be possible to obtain more of this super-archaic sequence by applying ARGweaver-D to the set of 161 Oceanian genomes recently sequenced [25], looking for super-archaic segments passed through the Denisovans.

There have been several studies suggesting super-archaic introgression into various African poulations [9, 10, 26]. However, ARGweaver-D only detected a small amount of Sup→Afr introgression, which was somewhat lower than our estimated false positive rate. One aspect to note here is that the power to identify introgression from an unsequenced population is highly dependent on the population size of the recipient population. The larger the population, the deeper the coalescences are within that population, making it more difficult to discern which long branches might be explained by super-archaic introgression. In the case of Africans, we used a population size of 23,700, which was our best estimate from previous runs of GPhoCS [7, 12]. If we had used a smaller population size, ARGweaver-D would have produced more Sup→Afr predictions, but most of these would be false positives unless that smaller population size is closer to the truth. Overall, we caution that the problem of detecting super-archaic introgression into a large and structured population such as Africas is very difficult, and that claims of such introgression need to be robust to the demographic model used in analysis. It may not be possible to address the question of ancient introgression into Africans without directly sequencing fossils from the introgressing population.

We also explored some introgression events that do not have any support from previous literature; namely the Hum→Den and Sup→Nea events. *A priori*, we expected that levels predicted for these events would likely serve to confirm our false positive rates in real data. However, it is also possible that there is some amount of these types of gene flow, which has not been detected previously because it goes against the net direction of gene flow. For the Hum→Den event, we predicted a slightly smaller fraction (0.37%) than our predicted false positive rate from simulations (0.41%). For Sup→Nea, we predicted 0.75% of the genome introgressed, which is slightly higher than our predicted false positive rate for this event (0.65%). While these fractions are small, it seems entirely plausible that if there was admixture between *Homo erectus* and the Denisovans, there may have also been some with Neanderthals, perhaps in the Middle East; or genes may have passed from *Homo erectus* to Neanderthal through the Denisovans. Given the number of known interactions between ancient hominins, it may be more reasonable to assume that gene exchange likely occurred whenever these groups overlapped in time and space.

## Materials and Methods

### General ARGweaver-D settings

For all ARGweaver-D runs in this paper, the MCMC chain was run for 2000 iterations, with the first 500 discarded as burnin, and ARGs sampled every 20 steps thereafter. Except where otherwise noted, phase was randomized for all individuals and the phase integration feature of ARGweaver-D was used (--unphased). We used site compression throughout (--compress 10). We also used --start-mig 100, which disallows migrations for the first 100 iterations of the sampler, enabling ARGweaver-D to establish an ARG with a good general structure before exploring the migration space.

#### Recombination rate

Rather than use a recombination map calculated from modern humans, which may not be accurate for ancient hominins, we used a constant recombination rate of 5e-9/bp/generation for all analyses. This value was chosen for being somewhat between the mean and median genome-wide recombination rates (1.3e-8 and 1.7e-9 per bp per generation, respectively), and for providing reasonable power while still maintaining a low false positive rates in simulations (see Supplementary Text). Note that all simulated data sets were nonetheless created with a real human recombination map (see “Simulations”, below).

#### Mutation rate

For real data analysis, the mutation rate map was based on primate divergence levels in 100kb sliding windows, using genome-wide alignments of human, chimp, gorilla, orangutan, and gibbon sequences (see Supplementary Text for details), and scaled to an average rate of 1.45e-8/generation/site. Simulated data sets were generated by sampling rates from this map, and the same map was used for analysis.

### Demographic model

The demographic model used in all ARGweaver-D analyses is depicted in Fig 3. The divergence times used were taken from [8], and population sizes from [7] (which were based on estimates from GPHoCS [12]). When analyzing chrX, population sizes were scaled by a factor of 0.75. When analyzing non-African humans, we only included the “recent” migration bands from Neanderthals and Denisovans into humans, whereas when looking for older introgression events, we excluded the “recent” bands as well as non-African humans.

Recall that ARGweaver uses a discrete-time model; 20 discrete times were chosen to span the range of relevant times, with more density near the leaves (where more coalescences occur) and to allow for coalescences between migration and population divergence events in the models. The discrete times (in kya) were: 0, 100, 200, 300, 400, 450, 500, 550, 600, 700, 950, 1200, 1450, 1700, 2000, 3000, 5000, 7000, 13,000, 15,000. Migration events occurred at half-time points including 50, 150, 250, and 350kya. Note that on this time scale, the European/African split is very recent, so that we did not model the population divergence among modern humans or recent growth in out-of-Africa populations. Similarly, we did not model the divergence between the Altai and Vindija Neanderthals, which are estimated to split only ∼ 15ky before the Altai Neanderthal individual lived. Throughout, we assume a generation time of 29 years [27].

### Calling introgressed regions

Once ARGweaver-D has been run, introgressed tracts can be identified for each migration event by scanning the resulting ARGs for local trees whose branches follow that migration band. Throughout this paper we use a probability threshold of 0.5 to identify introgressed regions, indicating that the region was introgressed in at least half of the sampled ARGs. To predict introgressed regions for a particular individual, we compute the posterior probability that either of the individual’s two haploid lineages are introgressed. The probability of being in a heterozygous or homozygous introgressed state can be calculated as the fraction of ARG samples in which one or two lineages (respectively) from an individual are introgressed in the local tree.

The coverage of introgressed regions for an individual is computed as one-half times the coverage of heterozygous regions, plus the coverage of homozygous regions. In theory, this fails to account for sites that switch between the heterozygous and homozygous states without reaching the threshold for either, but in practice this occurs at a negligible fraction of sites.

### Analysis of hominin data

#### Data preparation

We ran a series of ARGweaver-D analyses on freely available hominin data, described in Table 2. The panTro4 chimpanzee sequence was used as a haploid outgroup. The chimp alignment to hg19 was extracted from the alignments of 99 vertebrates with human available on the UCSC Genome Browser (http://hgdownload.soe.ucsc.edu/goldenPath/hg19/multiz100way). Any region which did not have an alignment for chimp is masked in the chimp sequence.

**Table 2.** Hominin samples used in this study.

| name | region | source | ID | sex | coverage | age (ky) |
| --- | --- | --- | --- | --- | --- | --- |
| Vindija Neanderthal | Europe | Max Planck | Vindija33.19 | F | 30x | 52 |
| Altai Neanderthal | Siberia | Max Planck | Altai | F | 52x | 115 |
| Denisovan | Siberia | Max Planck | Denisova | F | 31x | 72 |
| Papuan | Oceania | SGDP | LP6005441-DNA_B10 | F | 41x | 0 |
| French Basque | Europe | SGDP | LP6005441-DNA_D02 | F | 36x | 0 |
| Khomani San | Africa | SGDP | LP6005677-DNA_D03 | F | 44x | 0 |
| Mandenka | Africa | SGDP | LP6005441-DNA_F07 | F | 37x | 0 |

We downloaded samples generated by investigators at the Max Planck Institute from: http://cdna.eva.mpg.de/neandertal/Vindija/VCF; this directory contains genotype calls for several ancient genomes using a consistent pipeline and genotype caller (snpAD) throughout. SGDP: Simons Genome Diversity Panel.

#### Filtering

For each individual, we masked genotypes with quality scores less than 20 or sequencing depths outside the range [20, 80]. For each ancient individual, we also used the filters recommended by [8] and provided here: http://cdna.eva.mpg.de/neandertal/Vindija/FilterBed. We also masked (for all individuals): any site which belongs to a non-unique 35mer, according to the UCSC Genome Browser table hg19.wgEncodeDukeMapabilityUniqueness35bp; “black-listed” sites falling under the tables hg19.wgEncodeDacMapabilityConsensusExcludable or hg19.wgEncodeDukeMapabilityRegionsExcludable; ∼ 9% of the genome for which SGDP genotype calls were not provided (85% of this set overlapped previously mentioned filters). ARGweaver-D was run in 2.2Mb windows, but we excluded any window for which any of the ancient filters, or the combined site filters, exceeds 50% of bases. In total we analyzed 1,166 autosomal windows and 52 windows on the X chromosome, covering 2.56Gb of hg19.

#### CRF calls

Introgression calls from [5] were downloaded from https://sriramlab.cass.idre. ucla.edu/public/sankararaman.curbio.2016/summaries.tgz. As recommended by the README contained therein, “set1” calls were used for Neanderthal ancestry in the Basque individual, whereas “set2” calls were used for Denisovan ancestry in both the Basque and Papuan, as well as for Neanderthal in the Papuan. For each individual, we took the set of regions with probability of introgression *≥* 0.5 in either haplotype.

#### F4 Ratio

The F4 ratio statistic F4(Altai, chimp; Basque, African)/F4(Altai, chimp; Vindija, African) was calculated across the autosomal genome, and for each individual chromosome. For the African samples, we used allele frequencies across 29 African individuals from the SGDP data set (this excludes 15 individuals with the highest Neanderthal ancestry according to [11]). For this analysis we masked all sites that did not have a filter level (FL) field of 9 in the SGDP individuals. For the Neanderthals we used the same filters described previously.

### Simulated data sets (deep introgression)

We performed a series of simulations to assess ARGweaver-D’s ability to detect older migration events. Each simulated data set consists of a 2Mb region with 5 unphased diploid individuals and one haploid outgroup, mimicking the demographic histories and sampling dates of the individuals from the real data analysis. All simulations were produced with the software msprime [28].

The population tree used in the simulations is identical to the one depicted in Fig 3, and sampling dates correspond to the sample ages in Table 2. The human population size history also corresponds to the one in Fig 3. For the archaic hominins, we simulated a more detailed model of population size change, using piecewise-constant estimates produced by PSMC [29] and published in [8]. For the Neanderthal population history, we averaged the histories produced separately for the Altai and Vindija individuals, for the time periods when they overlap. Similarly, we averaged the Denisova and Neanderthal population size estimates during the time frame of their common ancestral population (415-575 kya).

For each data set, a random 2Mb region of the autosomal genome was chosen as a template region from which we chose recombination rates and mutation rates used to generate the simulated data. We used the recombination map estimated from African-American samples [30]. For the mutation map, we used the same map as in the real data analysis (based on primate divergence levels). Missing data patterns were also taken from the template region; we applied the same ancient genome masks and mapability/blacklist masks to the simulated data. (We did not mimic the sequencing depth or quality score masks, which affected a relatively small fraction of sites).

Overall we produced several sets of simulations, each consisting of 100 2Mb regions. One set served as a control and contained no migration events. All other sets each had three types of migration (Hum→Nea at a rate of 8%, Sup→Den at 4%, and Sup→Afr at 0.5%). The rates of each event were chosen so as to have enough events per data set to be able to assess power, while still being less common than the non-migrant state. They were also chosen (by trial and error) to produce roughly similar levels of predicted introgression as observed in the real data. The simulated data sets varied in the demographic parameters used (migration time and super-archaic divergence time). A smaller set of additional simulations was produced with population sizes scaled by 0.75 to see how power might change on the X chromosome (see Fig S5).

All false positive and true positive rates were calculated basewise; separate false positive and true positive rates were calculated for each type of migration in the ARGweaver-D model. To be classified as a true positive, the method must infer the correct type of migration in the correct individual. False positives presented here were assessed using the simulated data set with no migration.

### Simulated data sets (Nea**→**Hum introgression)

We also did a smaller simulation study to assess performance on the Nea→Hum event and compare performance to the CRF. Most of the settings were the same as above, except that we sampled 86 haploid African lineages and 4 Europeans, along with the two diploid Neanderthals and a haploid chimpanzee outgroup. Demographic parameters were the same as above, except that a European population diverged from the African population 100kya and had a initial size of 2100; at 42kya it experienced exponential growth at a rate of 0.002, for a present-day population size of 37236. (These parameters were roughly adapted from [31], but modified to reflect current smaller estimates of the mutation rate in humans.) We then added 2% migration from Neanderthal into Europeans at 50kya. In some supplementary analysis we also included 5% migration from human to Neanderthal 250kya.

For this analysis only, we used true haplotype phases, in order to have a fair comparison with CRF, which assumes phased samples.

#### Annotations

CDS, 3’UTR, and 5’UTR annotations were taken from the ensGene (ensembl) track on the UCSC genome browser. Enhancers and promoters were extracted from the Ensembl regulatory build dated 2018-09-25. PhastCons elements came from the phastConsElements46wayPrimates track on the UCSC Genome Browser.

## Acknowledgments

Thank you to Sriram Sankararaman for providing CRF software. MJH and AS were supported by US National Institutes of Health grant R35-GM127070 (to AS), and MJH was additionally supported by National Science Foundation GRFP DGE-1650441. ALW was supported by an Alfred P. Sloan Research Fellowship and a seed grant from Nancy and Peter Meinig. This work used the Extreme Science and Engineering Discovery Environment (XSEDE), which is supported by National Science Foundation grant number ACI-1548562. The content is solely the responsibility of the authors and does not necessarily represent the official views of the US National Institutes of Health.

## Supporting information

**Table S1.** Sup→Den regions overlapping Den→Hum regions predicted by the CRF

| Location (hg19) | count | overlapping genes |
| --- | --- | --- |
| chr15:56880301-56943860 | 10 | RP11-1129I3.1, ZNF280D |
| chr5:35268551-35472820 | 9 | U3 |
| chr15:79936231-80045380 | 7 |  |
| chr2:183978771-184038340 | 7 | NUP35 |
| chr1:40622111-40751801 | 6 | RLF, RNU6-1237P, TMCO2, RP1-39G22.7, ZMPSTE24 |
| chr4:143486431-143606100 | 6 | INPP4B, RP11-223C24.1 |
| chr15:63493991-63599658 | 5 | RAB8B, APH1B |
| chr17:30992260-31232970 | 3 | MYO1D, RP11-220C2.1, Y_RNA, AC084809.2, AC084809.3 |
| chr5:74577091-74897550 | 3 | CTD-2235C13.2, HMGCR, COL4A3BP, CTD-2235C13.3, POLK, RNU7-175P, CTC-366B18.2 |
| chr20:18369011-18456230 | 3 | DZANK1, RNA5SP476, POLR3F, MIR3192 |
| chr2:104441221-104575299 | 2 | AC013727.1, AC013727.2, RP11-76I14.1 |
| chr3:156394341-156515810 | 2 | TIPARP, RP11-392A22.2 |
| chr8:97918711-98192640 | 2 | CPQ, KB-1958F4.2, KB-1958F4.1 |
| chr8:56673021-56798570 | 2 | TMEM68, TGS1, LYN |
| chr13:77606499-77899730 | 1 | MYCBP2, MYCBP2-AS1, RP11-226E21.2 |
| chr4:85723291-85798820 | 1 | WDFY3, RP11-147K21.1 |
| chr6:131062559-131237440 | 1 | SMLR1, EPB41L2 |
| chr7:83335141-83452959 | 1 |  |
| chr10:52582401-52700350 | 1 | A1CF, RP11-449O16.2 |
| chr3:129951121-130100515 | 1 | COL6A5, AC093004.1 |

The “count” column shows the number of non-African SGDP individuals who have Denisovan introgression at this locus. We restricted this list to Sup→Den regions for which at least 90% of SGDP individuals without Denisovan introgression have a higher divergence to the Denisovan than to Neanderthals.

**Table S2.**
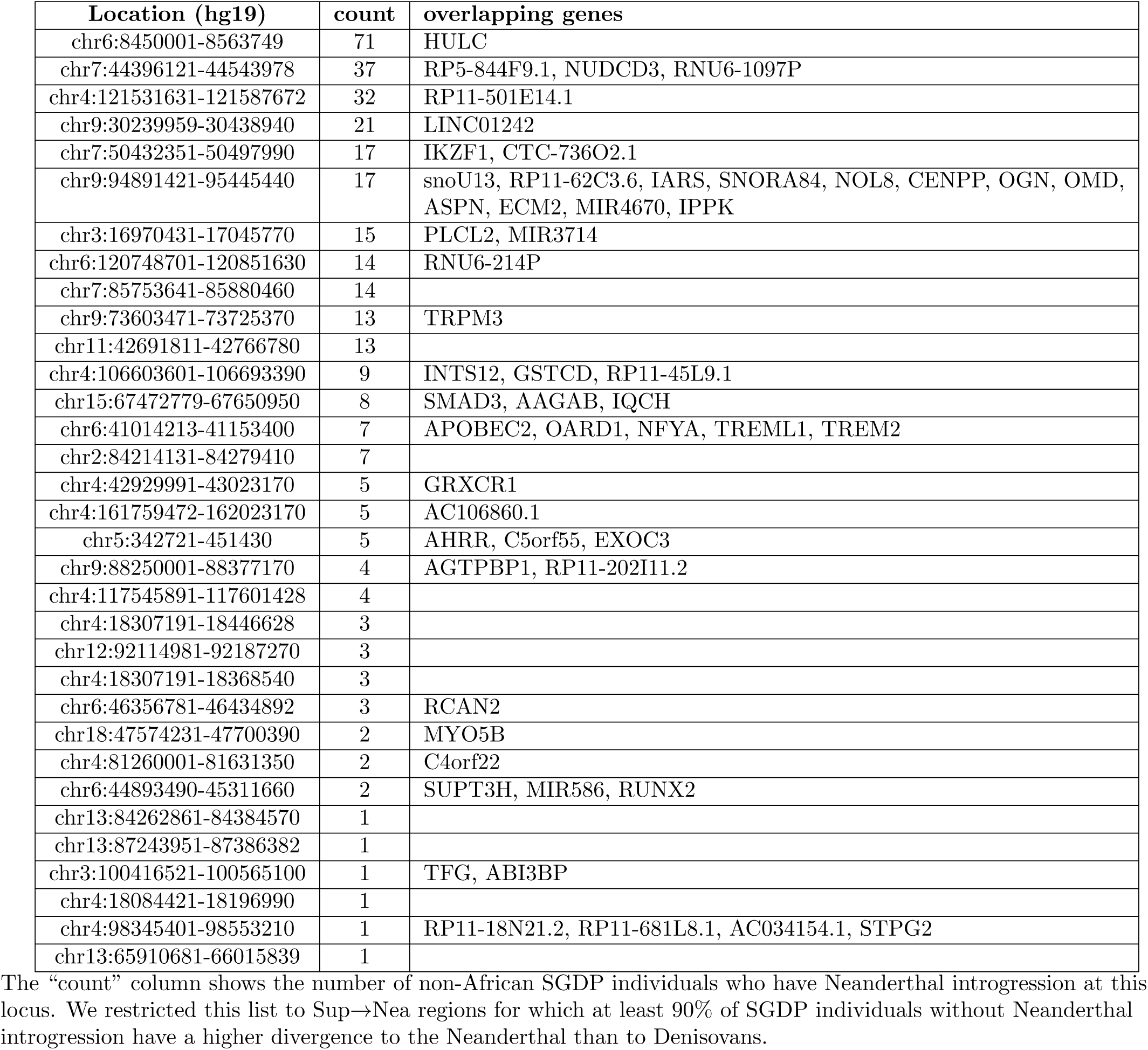
Sup→Nea regions overlapping Nea→Hum regions predicted by the CRF

**Fig S1.**
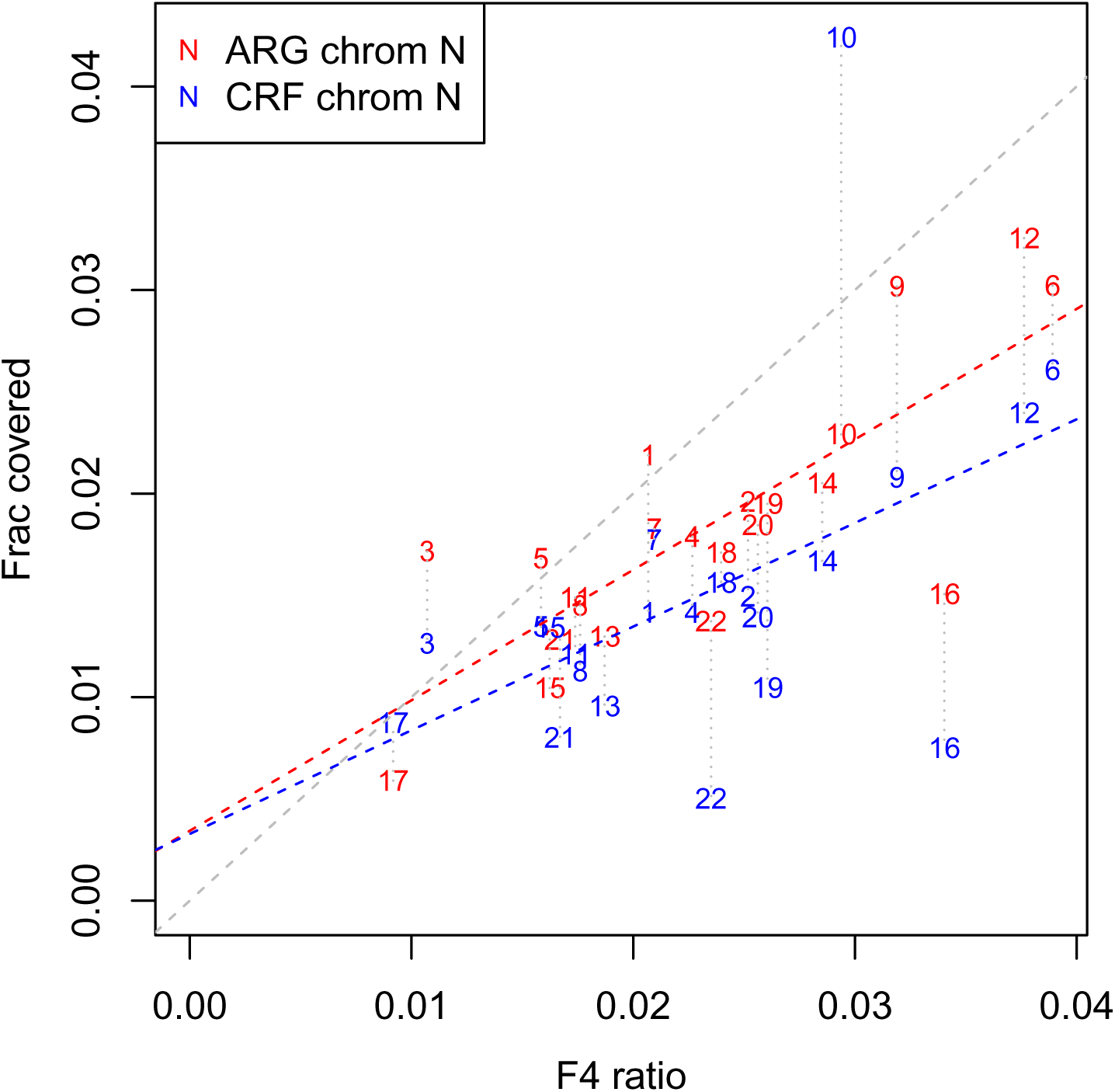
Coverage of introgression predictions vs. F4 Ratio. For each autosomal chromosome, we plot the expected fraction predicted introgressed by ARGweaver-D (red) and CRF (blue) into the Basque individual, vs the F4Ratio statistic F4(Altai, chimp, Basque, Afr)/F4(Altai, chimp, Vindija, Africa) computed for variants on each chromosome. The dashed red and blue lines show the best linear fit for each method, whereas the gray dashed line shows *x* = *y* for reference. Dotted gray lines connect results from the same chromosome.

**Fig S2.**
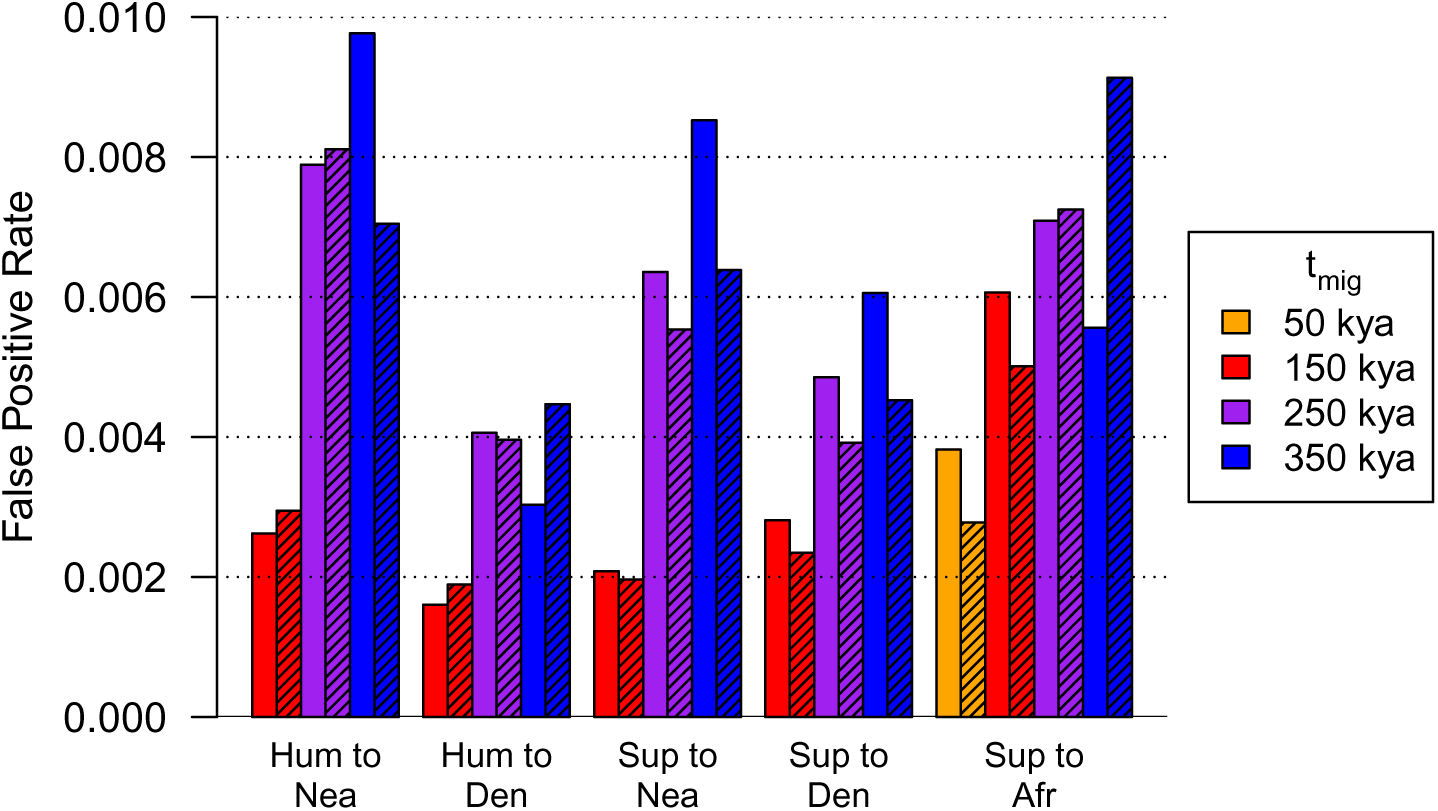
False positive rates calculated from simulations, using several ARGweaver-D models. Color indicates the value of *t_mig_*. Shaded bars have *t_div_* = 1.5Mya and solid bars have *t_div_* = 1.0Mya. False positive rates are calculated base-wise using a posterior probability cut-off of 0.5. The same set of underlying data was used for all the calculations in this plot; it was simulated as in Fig 3, but with no true migration events.

**Fig S3.**
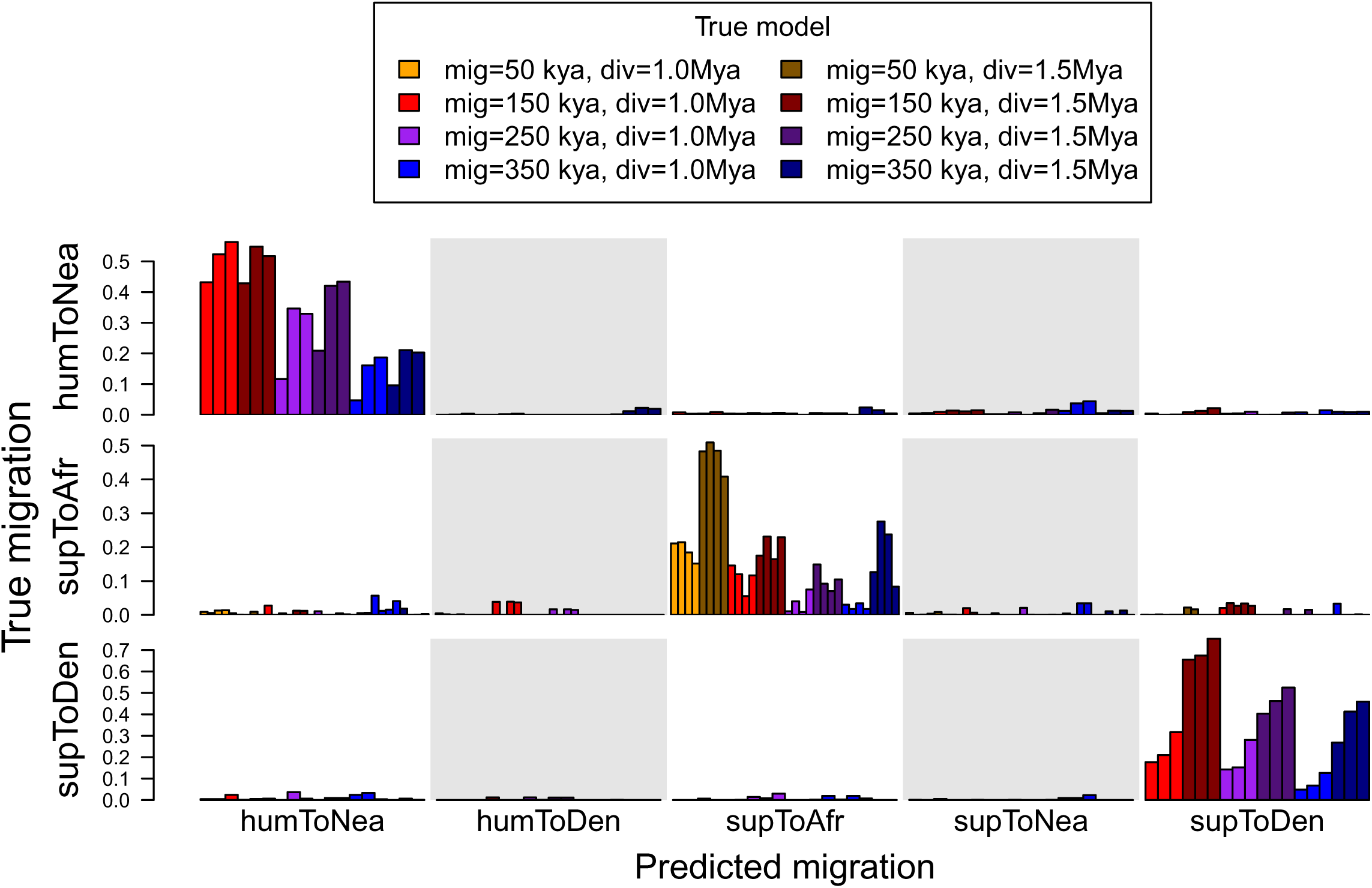
Detailed simulation results. Each row shows a true migration category, and each column shows the fraction of bases predicted in the category indicated at the foot of the column. The color of each bar represents the true parameters used in simulation, as indicated in the legend, with darker colors used for the older super-archaic divergence time. Multiple bars of the same color show results on the same data set, using an ARGweaver-D model with a different *t_mig_*. The value of *t_mig_* used by ARGweaver-D is not indicated in the plot, but increases from left-to-right: *t_mig_* = 50, 150, 250, 350kya, with 50kya only shown for Sup→Afr. All the models used *t_div_* = 1Mya; the plot with *t_div_* = 1.5Mya is nearly identical.

**Fig S4.**
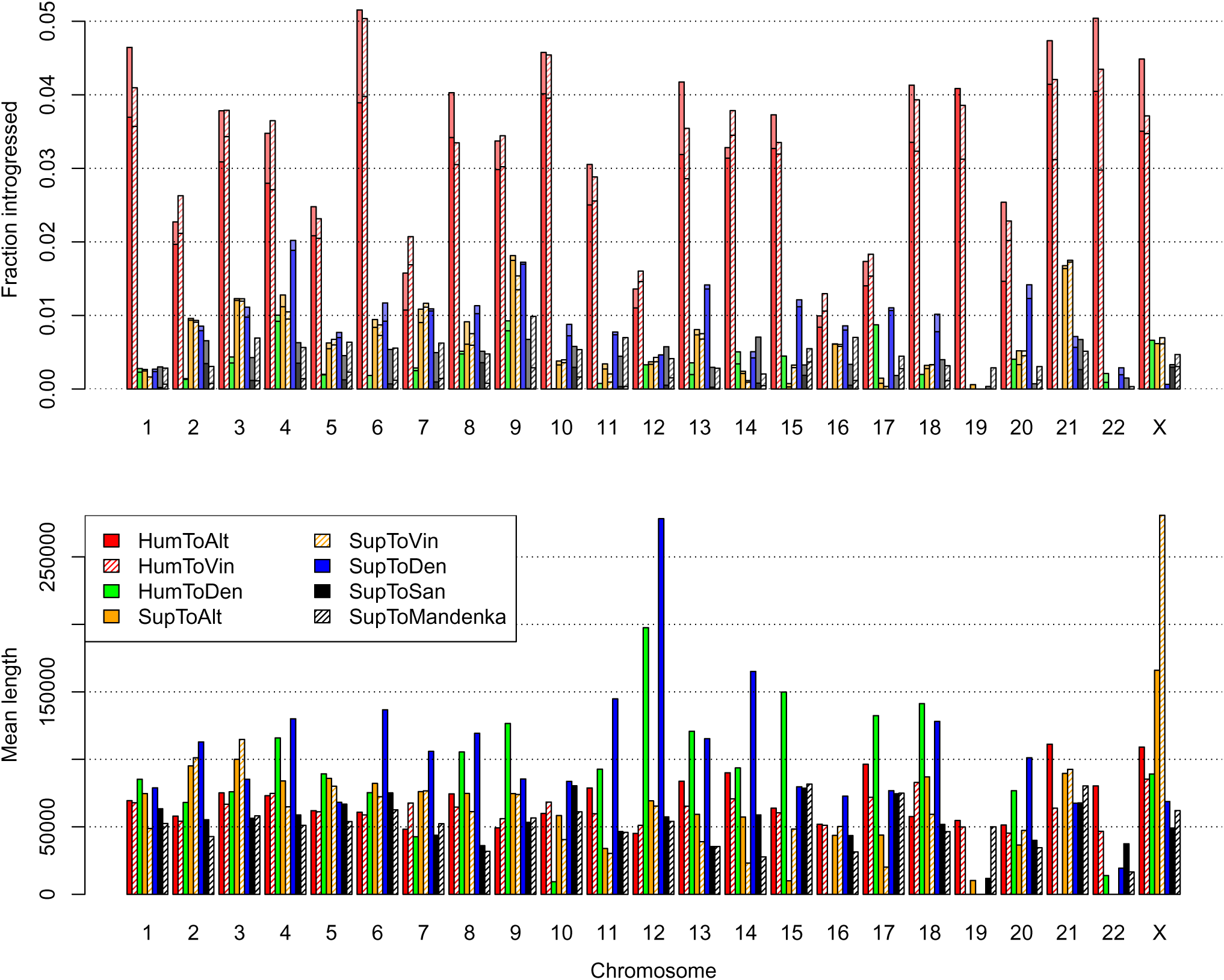
Properties of introgressed regions by chromosome. The top plot shows average coverage of predicted introgressed regions per haploid genome, with darker portions representing homozygous regions. The bottom shows average length of introgressed regions by chromosome.

**Fig S5.**
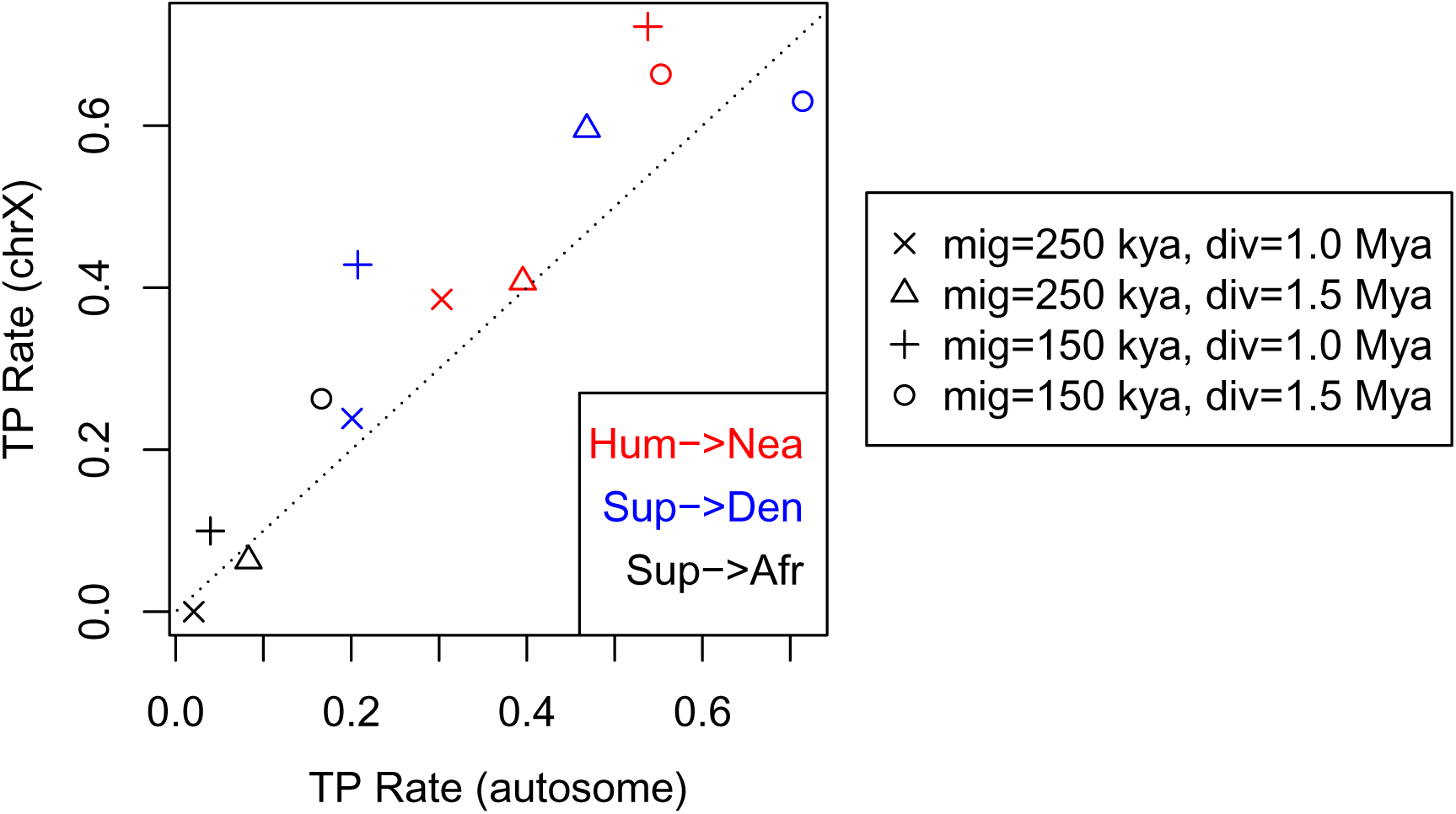
True positive rate for simulations on X chromosome vs autosomes. The y-axis shows true positive rates from simulations where population sizes were multiplied by 0.75 to roughly approximate X chromosome demography. Different plotting characters are used for different simulation models, as indicated in the legend. All ARGweaver-D analysis was done with *t_mig_*=250kya and *t_div_*=1.0Mya.

**Fig S6.**
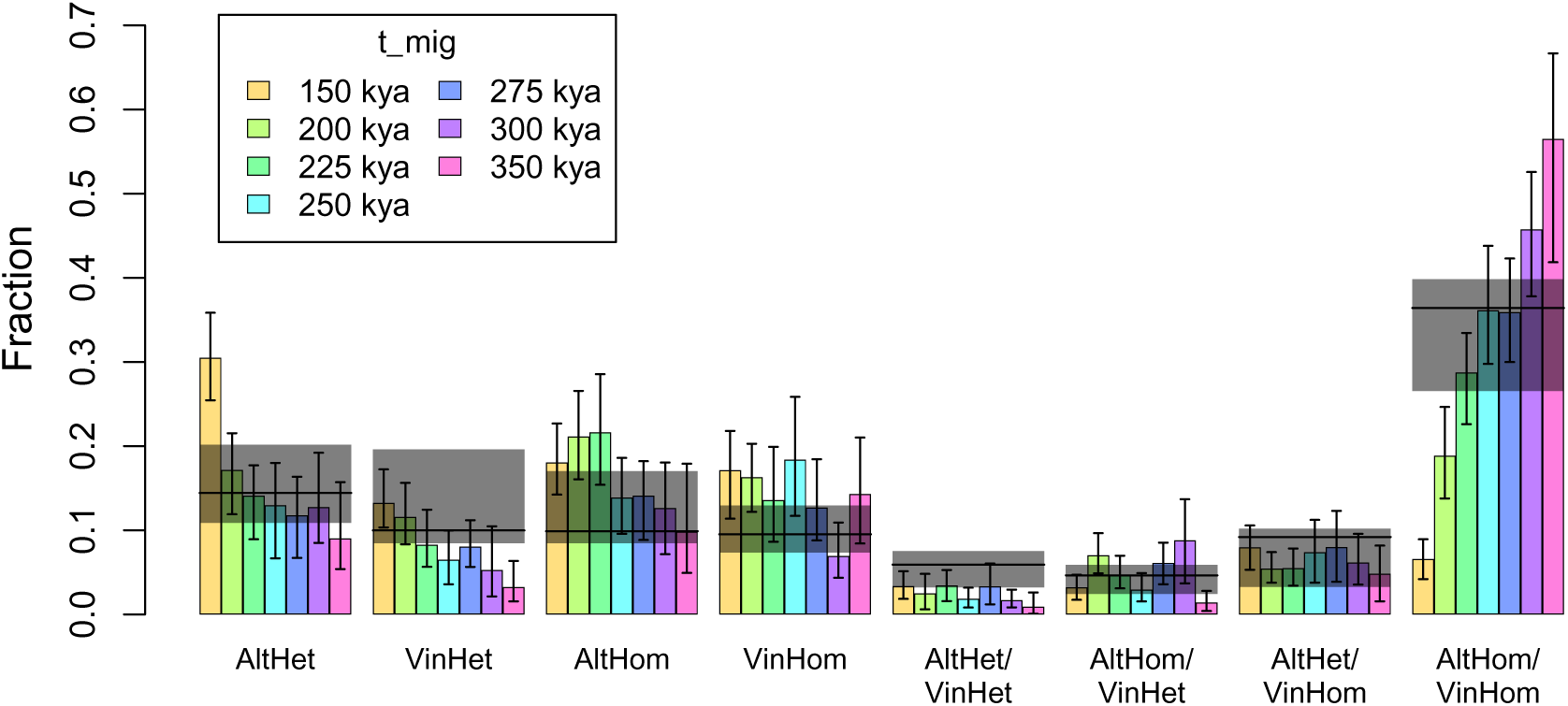
Frequencies of Hum→Nea introgression categories. For both the real and simulated data, Hum→Nea regions were ascertained with ARGweaver-D using a model with *t_mig_* = 250kya and *t_div_* = 1Mya. These regions were classified as heterozygous/homozyogus in the Altai Neanderthal (AltHet/AltHom), and in the Vindija Neanderthal (VinHet/VinHom), depending on which branches are in the migrant state in the majority of sampled ARGs. Here, the colored bars represent the fraction of Hum→Nea bases in each category for simulated data sets generated with different values of *t_mig_*; the error bars show 95% confidence intervals (CIs) computed using 100 bootstrap replicates across the introgressed elements. The horizontal black lines represent the amount observed in the real data, with the gray boxes showing the CIs, also obtained by the same bootstrap process.

**Fig S7.**
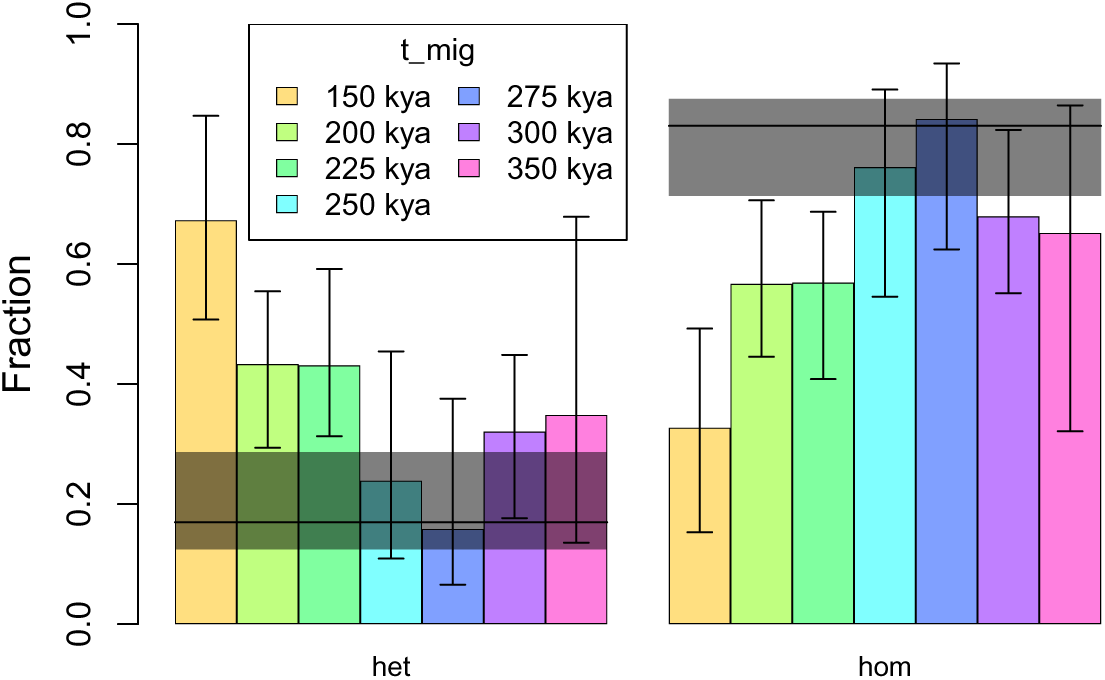
Frequencies of Sup→Den introgression categories. This figure is analygous to Fig S6; here we look at putative Sup→Den regions. Because there is only one Denisovan individual, there are only two categories: heterozygous or homozygous. Note that while we expect rates of heterozygosity to decrease with migration time, the confidence intervals here are wide, and there may be conflicting ascertainment effects that cause the apparent increase in heterozygous segments for the simulated data sets with oldest *t_mig_* values.

**Fig S8.**
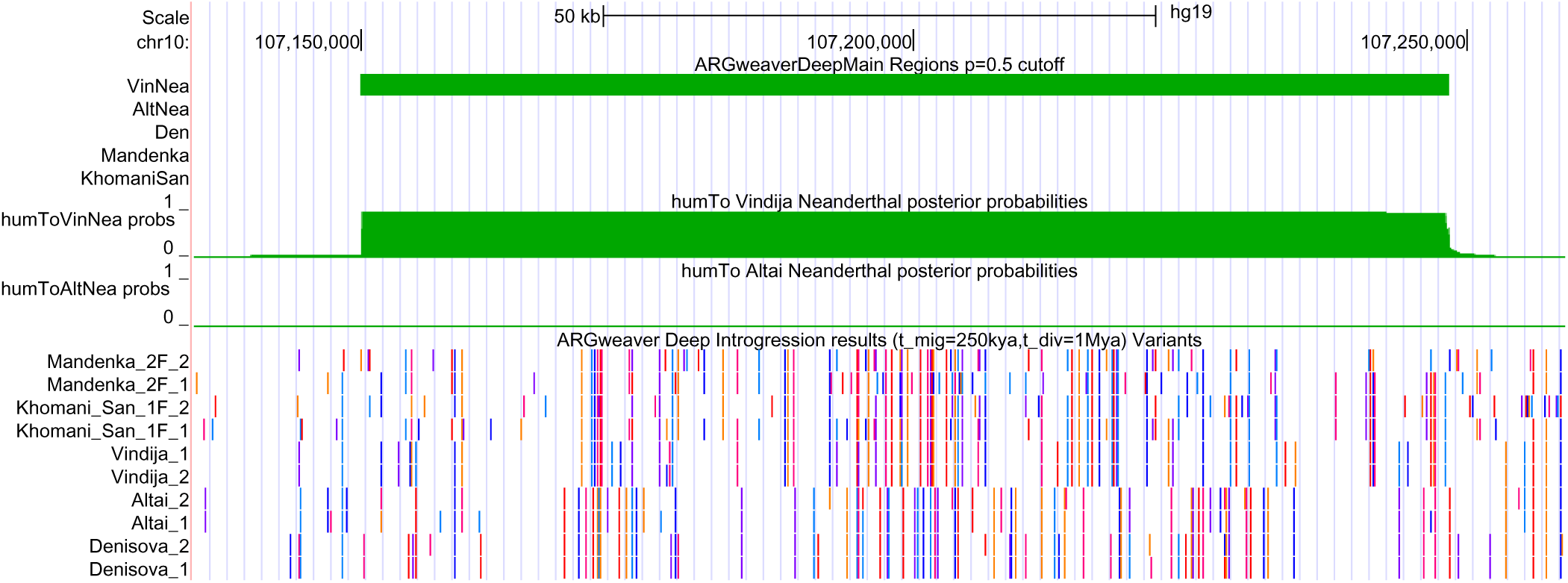
UCSC Genome Browser shot of a region with predicted homozygous Hum→Nea introgression in Vindija. This region on chromosome 10 has a high-probability introgressed region in both Vindija (but neither Altai) haplotypes. The top green bar indicates a predicted Hum→Nea region in Vindija, and below this is the posterior probability of introgression across the region in both Neanderthals. The variant track is similar to Fig 8. Here, we see almost identical haplotypes between Vindija and the Africans, whereas Altai shares haplotypes with the Denisovan.

**Fig S9.**
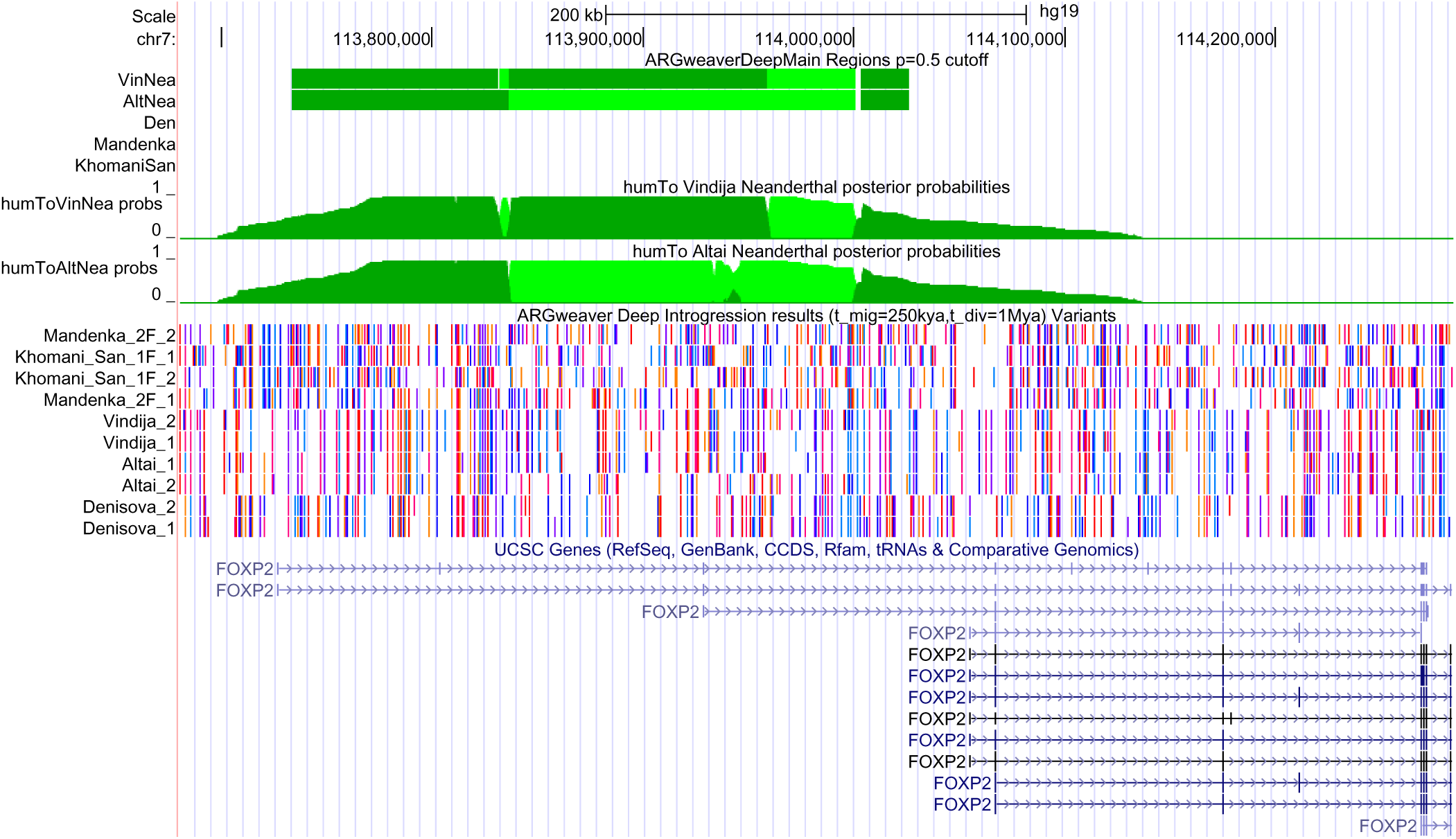
UCSC Genome Browser shot of a predicted Hum→Nea region overlapping FOXP2. Exon 7, which contains human-chimp substitutions shared by Neanderthals that may be involved with human speech, is located at the very right of this plot, and is not predicted introgressed. As in Fig 7, the light green implies heterozygous Hum→Nea introgression, whereas dark green is homozygous.

**Fig S10.**
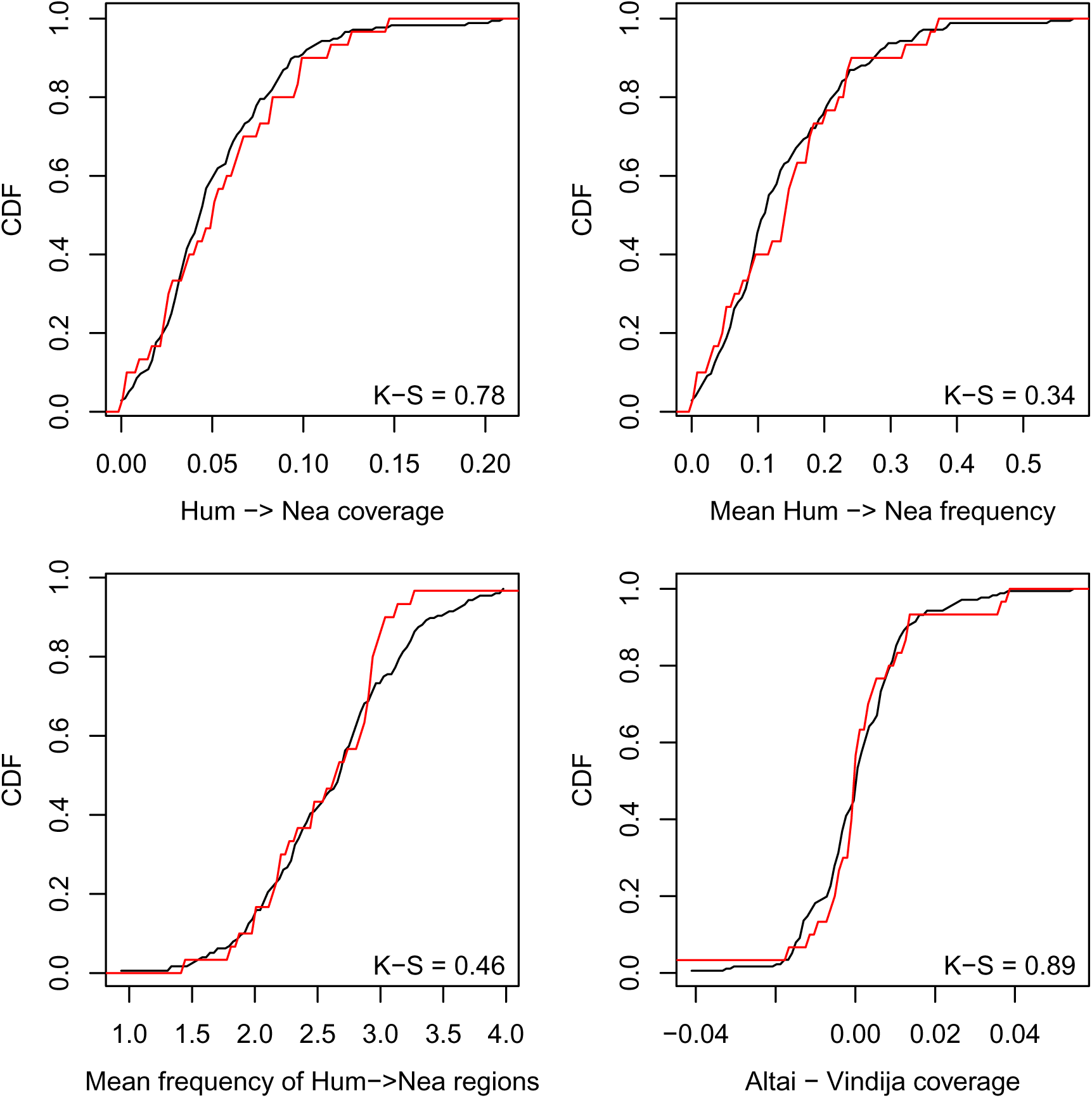
Properties of Hum→Nea regions within Nea→Hum deserts. We compared the distribution of various statistics across all non-overlapping 15Mb windows in the genome (black), to the distribution within deserts of Neanderthal introgression in humans of at least 10Mb (red). We excluded any window that crosses a telomere or centromere, or where ≥ 50% of the window does not pass our filters. In the bottom-right corner of each plot is shown the Kolmogorov-Smirnov statistic p-value, indicating that there is no significant difference between the black and red distributions. The statistics shown are indicated on the x-axis label. “Hum→Nea coverage” is average fraction of the window that contains any Hum→Nea region. “Mean Hum→Nea frequency” is the average number of introgressed haploid lineages of Hum→Nea across the window (where a frequency of zero indicates no introgression, and a frequency of 4 indicates homozygous introgression in Altai and Vindija). “Mean frequency of Hum→Nea regions” is the mean frequency, among regions with Hum→Nea calls. “Altai - Vindija coverage” is difference in mean coverage between the Altai and Vindija within each window.

