## Supplemental Text for "Mapping gene flow between ancient hominins through demography-aware inference of the ancestral recombination graph"

### Supplementary Methods

#### Threading an ARG conditional on population structure

The main engine behind ARGweaver-D is the "threading" operation, which samples the coalescence points for a lineage that has been removed from the ARG. The threading operation uses an HMM in which the state space changes along the genome as the local tree changes. At a given genomic location, the state space is defined as the set of all possible coalescence points on the local tree, given by every time point along every branch. In the multiple population model, the state space is augmented with a third dimension indicating the "population path" of the new branch. Each population path is a vector of population assignments at every time point. This population path is sampled as part of the threading procedure, and retained in the new ARG, so that the full ARG defines the local genealogies as well as the population assignments for every lineage at each time point.

Recall that in the original ARGweaver model, time is discretized into $K+1$ points, $t_0, ..., t_K$, with half-time points $t_{1/2}, ..., t_{K-1/2}$ between them. Coalescence and recombination events occur only at whole time points, based on their cumulative probabilities between the adjacent half-time points. In the multiple population model, both migration and population divergence are assumed to occur instantaneously at one of the discrete half-time points. This separates the coalescence process from the migration process, preventing ambiguities about the order of events in the ARG, and ensures that the number of lineages within a population is well-defined throughout a coalescence rounding interval.

Let each state be described by the vector $(b, t, p)$, where $b$ is a branch of the local tree, $t$ is a time (looking in backwards in time; $t = 0$ is present-day), and $p$ is a population path vector, such that $p_i$ gives the population of path $p$ at time point $i$. Each branch of the tree has its own population path, $p^{(b)}$. A state is only valid if $p_t = p_t^{(b)}$, since coalescence can only occur if the lineages are in the same population at the same time. We also assume throughout that each sample comes from a single known population (although this model could easily be extended to work for unknown or admixed samples). Therefore, for leaf branches, $p_0$ is fixed (or for an ancient sample with date $a$, $p_a$ is fixed).

Due to these constraints, only a subset of all population paths are valid for each possible coalescent point. In the absence of migration bands, every coalescence point on the tree will be reachable by either zero or one population paths. Therefore, the size of the state space will be reduced compared to the single population model. However, as migration bands are added, more coalescence points become reachable, and some will be reachable by multiple distinct population paths. The result is that the size of the state space - and the computational complexity of the HMM - increases as more migration bands are added, and can quickly exceed the single-population case. The cost of each migration band, in terms of run-time, depends on how many samples are in the receiving population, as well as the time of the migration band (recent migration events increase the state space more than older ones in a particular population). In order to improve both efficiency and mixing of the MCMC chain, we allow at most one migration event in any local tree in the ARG. So, ”double migration” or ”back migration” events are not allowed. When threading a branch into a local tree that already contains a migration, only states with non-migrating paths are valid. This assumption is reasonable when the rate of migration is low and the number of samples is modest.

Given this multiple population model, the threading algorithm proceeds similarly to the one described in the original ARGweaver paper [1]. The emissions probabilities (computed as the probability of the sequence data conditional on the local tree) are not affected by this model; nor are the probabilities of recombination at any point in the local tree. However, the probability of coalescence is now calculated conditional on the population path. The probability of the path also needs to be taken into account (as the product of migrating or not migrating as the branch passes through migration bands). Additionally, the symmetries exploited by ARGweaver for optimizing the forward algorithm also change. The algorithm takes advantage of the fact that the transition probabilities from state $x_{i-1}$ to state $x_i$ are not dependent on the branches assigned to each state, except for when the two branches are equal. Otherwise, only the coalescent times for each state matter, and the forward algorithm can be performed in $O(LnK^2)$ time, where $L$ is the number of sites, $n$ is the number of samples, and $K$ the number of time points. In the demographic-aware model, the calculation also depends on the population path assigned to each state, so the complexity increaes to $O(LnK^2P^2)$ where $P$ describes the maximum number of population paths that any lineage is able to take under the specified demographic model.

**Threading migrant lineages**

One additional difference is that a new type of threading has been implemented for the population model. The original model has both leaf threading (which removes and re-threads a single haploid lineage from the entire ARG), and subtree threading (which removes and resamples a series of branches, both internal and leaf). We found that neither of these algorithms are sufficient to achieve good mixing of the MCMC chain when old migration events are present, because they are not able to add or remove entire migrant haplotypes in one step.

To remedy this, we have added a branch removal algorithm that focuses on lineages and time points which may potentially reach a migrant state. Recall that subtree threading uses a ”branch graph” structure that is designed to choose a series of removal branches from adjacent local trees, so as to minimize constraints on how these branches need to be re-threaded to maintain consistency with the remaining ARG. Given a removal branch at site $i$, the choice of removal branch at site $i+1$ is often deterministic, as most branches have a single analog in the neighboring tree with the exact same set of descendants. But when a branch is involved in recombination and recoalescence, then there may be two possible analogous branches to choose from. The original subtree threading algorithm made this choice randomly, and also required a Metropolis-Hastings rejection step to correct for the differing numbers of possible un-threadings in different ARGs (which is related to the number of recombination events in each ARG).

In our modified threading algorithm, we start by randomly choosing a migration band and a haploid lineage from a population which may follow this band. (For example, we may have a migration band between the 4th and 5th time interval to represent human to Neanderthal introgression, and we would randomly choose one of the Neanderthal lineages in the tree.) At the first site, we choose the branch ancestral to our chosen lineage which crosses through the time of the migration band (whether it migrates or not). We then follow the branch graph procedure, with the additional constraint that we only choose branches which span the time of the migration. In this way, there is never a random choice to make; if a branch is split in two by recoalescence of another branch onto it, then only one of the resulting segments spans the time of the migration band. In some cases, the chosen branch may be broken by recombination, and recoalesce more recently than the migration band; in this case we go back to the default of choosing the ancestral branch of the chosen lineage which crosses the migration time.

This modified-subtree algorithm guarantees that any migration event undertaken by an ancestor to the chosen lineage is completely removed from the ARG, and helps the MCMC sampler move to likely migration states, and also prevents the MCMC chain from getting "stuck" in a migration state. Note that this procedure is agnostic to whether any migration events actually exist in the ARG, and that the choice of a lineage and migration band is independent of the current sampled ARG. Therefore, the Metropolis-Hastings acceptance ratio is always 1, and the rejection step is unnecessary.

The effect of this threading algorithm is demonstrated in Figure S1, which shows performance with and without the new algorithm on simulated data. The algorithm appears to make very little difference in detecting the Nea→Hum migration event. We expect this is because for recent migration events, the leaf threading algorithm can effectively remove and replace entire introgressed regions. However, the algorithm improves the power for the deep introgression events significantly (except for Sup→Afr, which has very little power in either case). When the algorithm is not used, the power is much lower, though the false positive rate is still low. In this example at least, then, there is not a problem with getting "stuck" in a migration state, but in moving to the migration state.

All other analyses presented in this paper use this new threading algorithm. By default, when a migration model is being used, ARGweaver-D uses leaf threading algorithm for half the iterations, and equally divides the other iterations between the original and modified subtree pruning algorithms.

#### Ancient sample ages

Whereas the original implementation of ARGweaver assumed that all samples came from present day, there is now an option to specify sample ages. The option `--age-file` takes a file with the ages of ancient samples, and was used throughout this study to model the Neanderthal and Denisovan lineages. All sample ages are rounded to the nearest time point in the ARGweaver-D model.

Implementing this option required a few minor modifications to the software, including removing the implicit assumption that the distance from leaf to root is the same for all leaves. An ancient sample with age $t$ has leaves that

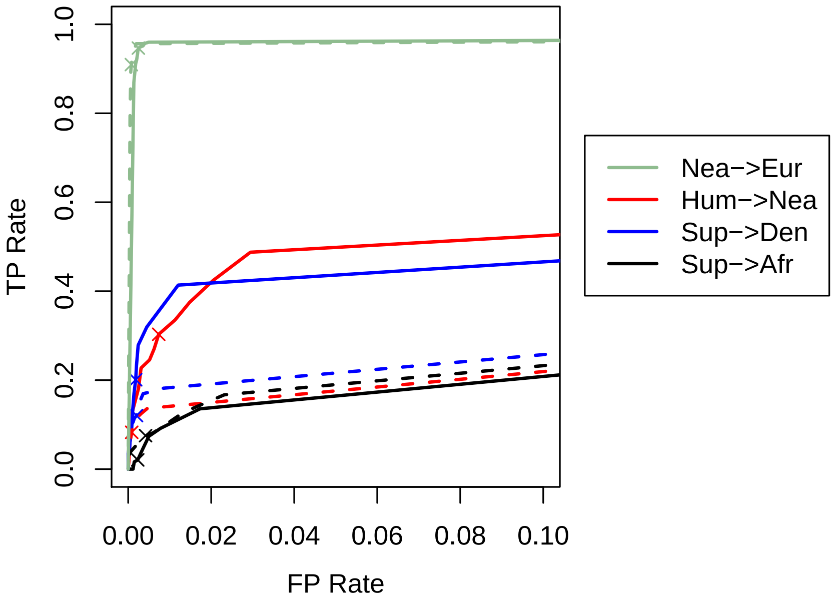


Figure S1: This ROC plot shows performance with (solid) and without (dashed) the modified subtree pruning algorithm. The Nea→Eur lines come from simulations in the main paper with a single migration at 50kya. The other lines come from "deep introgression" simulations with migration at 250kya and super-archaic divergence of 1Mya, and analyzed with the same model. The "x" on each line represents the performance when a posterior probability cutoff of 0.5 is used.

start at time $t$ instead of zero, and when threading this lineage, the only valid states are ones with ages $\geq t$. The code also had to be altered to ensure that lineages for an ancient branch do not contribute to coalescence probabilities in the time range $(0,t)$. Whereas the number of branches usually only decreases looking backwards in time, with ancient samples, branches may come into existence and the branch count can increase.

We did a simple simulation study to demonstrate that this option works as expected. Using `msprime` [2], we simulated 10 haploid lineages from a population of size 10000 across a 2Mb region. Four of the samples were modern day (sampled at $t = 0$); the other six were sampled at increasingly ancient times (50kya, 100kya, 150kya, 200kya, 250kya, 300kya). We used a recombination rate of 1.5e-8/recomb/bp/generation and a mutation rate of 2.5e-8/bp/generation. For this demonstration we treated samples as haploid with known phase. We then compared properties of ARGs inferred with the `--age-file` option, to ARGs inferred without the option, and also to the true known ARG. Figure S2 shows that this option effectively corrects for biases in the estimates caused by ignoring sample ages. It also shows that, even in this fairly extreme example with several very old samples, many statistics of the ARG are not too badly skewed even when the sampling ages are not properly taken into account.

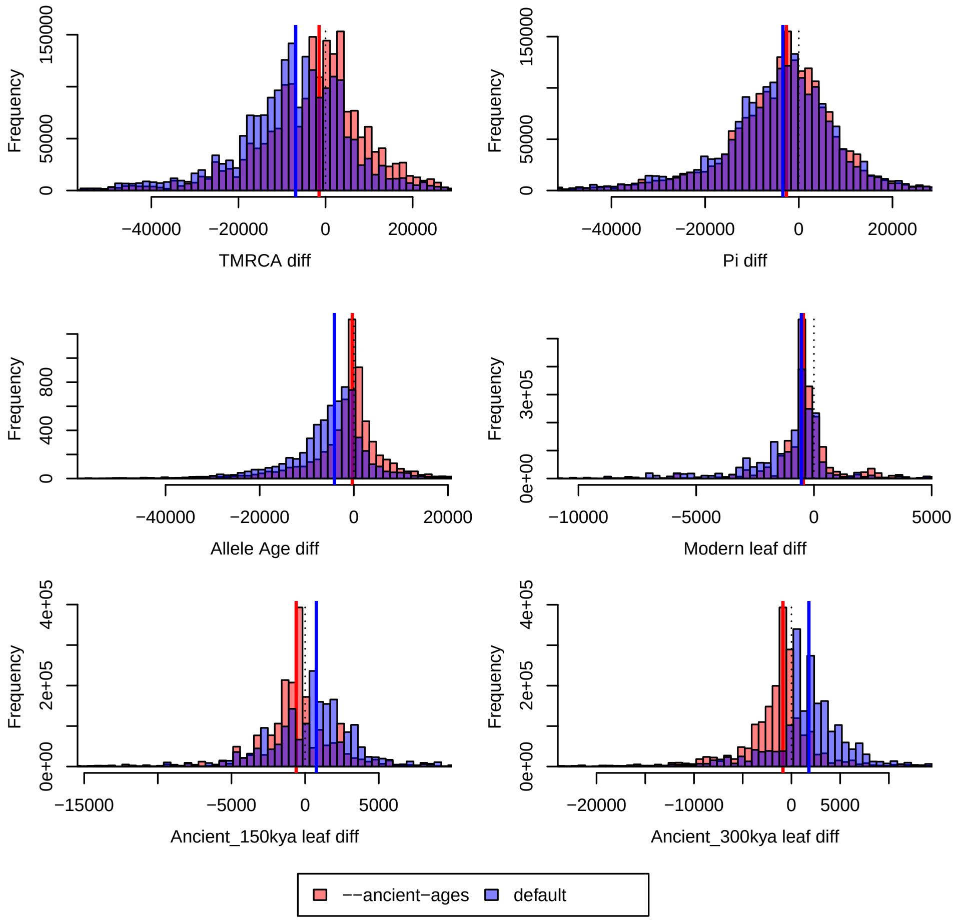


Figure S2: Looking at the ARG across a 2Mb region, at every base we compute the difference between the true statistic and the median from ARGs sampled across 2000 MCMC iterations. The pink distribution shows the ARGs inferred while accounting for ancient sampling dates; the blue uses the default parameter. (Purple is the overlap between the two). The dotted black line is at $x = 0$, and the red and blue lines are at the medians of the pink and blue distributions. The statistic for each plot is named in the x-axis, and the names are as follows: TMRCA (time to most recent common ancestor, in generations); Pi (average distance between two leaf nodes, in generations); Allele age (age of derived alleles); Modern leaf (coalescence time of a leaf node for a present-day sample); Ancient_150kya leaf (coalescence time of a leaf sampled 150kya), and Ancient_300kya leaf (coalescence time of a leaf sampled 300kya).

### Supplementary simulation results

#### Out-of-Africa simulations with Hum→Nea

In addition to the out-of-Africa simulations with Nea→Eur migration presented in the main paper, we performed a second set of simulations that also included a true Hum→Nea migration event at 250kya at a rate of 0.02. We ran both CRF and ARGweaver-D on this data set. We tried two different ARGweaver-D models, one with a single Nea→Eur migration band at 50kya, and one that also included a Hum→Nea band at 250kya. The introgression predictions are summarized in Figure S3. When ARGweaver-D has only one band, it performs similarly to CRF, though with a somewhat lower false positive rate. Both methods mis-classify a small fraction of Hum→Nea regions as Nea→Eur (9.0% for CRF and 7.6% for ARGweaver-D). When a second band is added to the ARGweaver-D model, the true positive rate (34%) for correctly identifying Hum→Nea is identical to what we saw in the ”deep introgression” simulations presented in the main paper. However, a large fraction (38%) of the Nea→Eur regions are incorrectly classified as Nea→Hum. We are not sure why the misspecification is so much higher in one direction than the other. However, we have observed in general that getting the correct directionality when there are migrations between sister populations is difficult. This is one reason for excluding non-African populations when looking for older migration events.

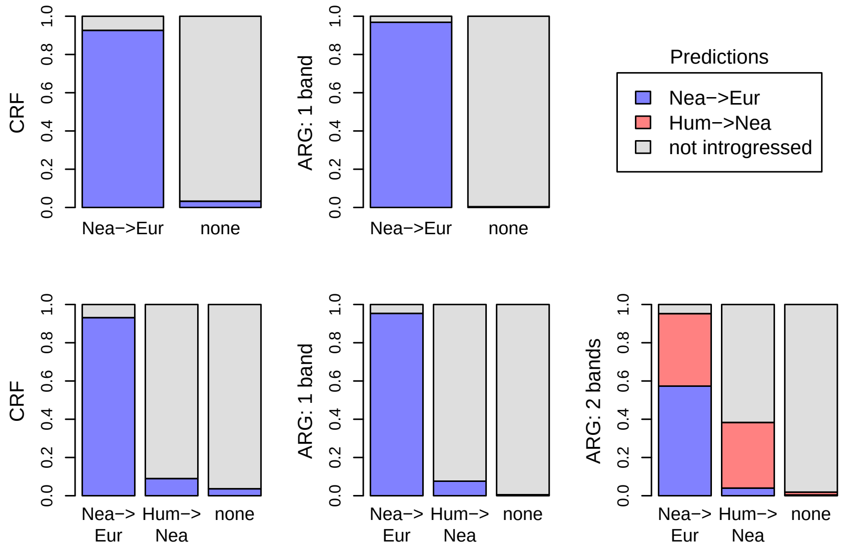


Figure S3: On the top is shown predictions for a set of simulations with only Nea→Hum introgression. Each bar represents a true (known) category; the colors show predictions for this category using the CRF (left) and ARGweaver-D with a single migration band (right). On the bottom are results where there are two true migration events. Here there are two sets of ARGweaver-D results; one with only the Nea→Hum band, and one that also has a Hum→Nea band.

###### Recombination rate analysis

Most of the analysis in this paper was done assuming a constant mutation rate across the genome. While this is unrealistic, little is known about the recombination map in Neanderthals and Denisovans (and nothing is known about possible super-archaic recombination maps).

To explore the effects of recombination rate misspecification, we compared the performance on our "deep introgression" simulations with four different recombination settings: one with the true map used in the simulations, one with a constant rate of 5e-9/recomb/generation/base-pair, another with a higher rate of 1e-8/recomb/generation/base-pair, and one with an incorrect recombination map. All simulations were created with a true recombination map sampled from some a random region of the human recombination map generated from African-American samples [3]; the incorrect maps were sampled from a different random region of the human genome. The results are shown in Figure S4.

Overall, the recombination rate seems to have the biggest impact on the false positive rate. Except for in Sup→Afr, where the power is low everywhere, the best performance is when the true recombination rate is used, and the worst are when a too-high or wrong rate is used. Using a constant rate of 5e-9 gave intermediate performance, and fewer false positive than other incorrect maps. Because hotspots identified in human may not apply to Neanderthal, Denisova, or super-archaic hominins, we chose to use the low recombination rate.

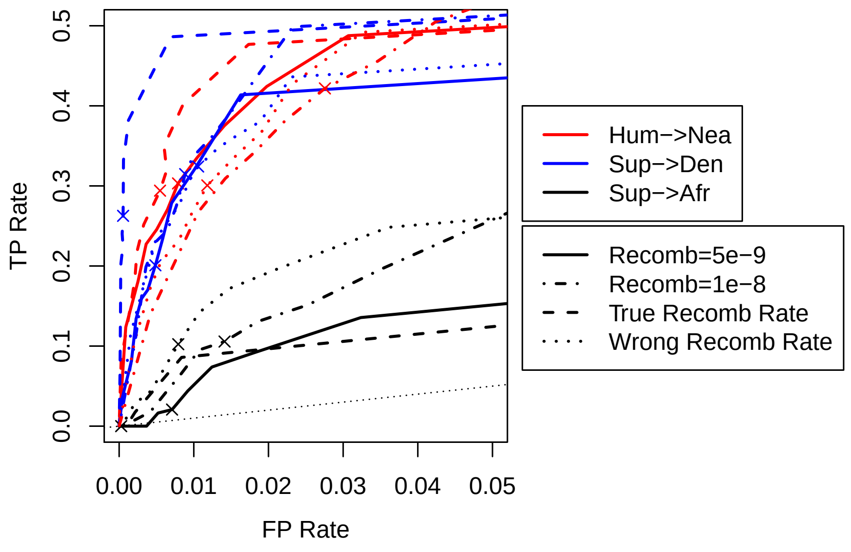


Figure S4: Performance on the "deep introgresion" simulations, in which the only difference between ARGweaver-D runs is the recombination map used for analysis.

###### Number of African samples

In theory, larger numbers of African samples might be expected to improve performance for finding Hum→Nea introgression events, as the introgressing ancient human population is most closely related to modern Africans, and

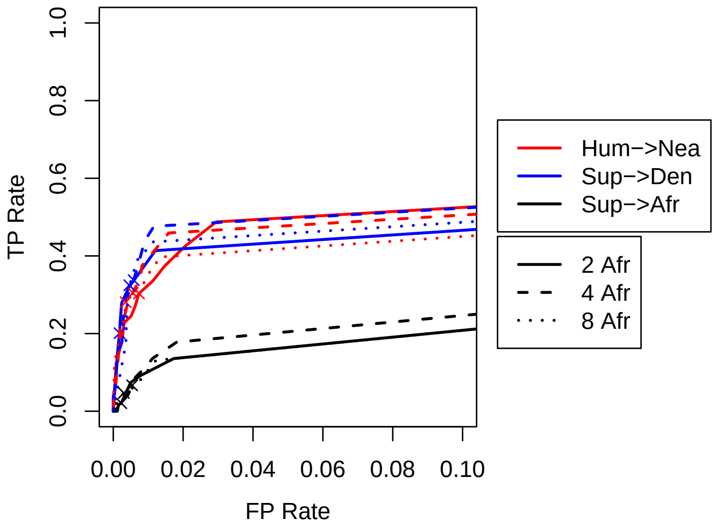


Figure S5: Effect of using more African individuals in the analysis. Here, we used the same "deep introgression" simulations as in the main paper, but sampled and used more African individuals in the ARGweaver-D analysis. Both the simulated data and the ARGweaver-D model had a migration band at 250kya and super-archaic divergence of 1Mya. The "x" on each line represents the performance when a posterior probability cutoff of 0.5 is used.

seems to be equally related to various diverged African populations [4]. It is also possible that more African samples could boost the power of detecting Sup→Den regions, as they contribute more information about the ancestral archaic population. However, in practice we observed that power using 2 Africans was similar to using 4 or even 8 African individuals (Figure S5). We are not sure why this is, but suspect that the MCMC sampler does not mix as well as more individuals are added. Because ARGweaver-D is faster with fewer individuals and it does not seem to have much effect on performance, we did our main analysis with two Africans.

### Supplementary analysis

#### Lengths of real vs simulated introgressed regions

The lengths of introgressed regions should be informative for the time of migration. However, there is more power to detect longer regions, creating a strong ascertainment bias that makes interpretation of the lengths difficult. The rate of recombination is also an important factor affecting the distribution of lengths, and the recombination rates in Neanderthal and Denisovan are not well characterized. Still, we looked at the distribution of lengths in our predicted set, and compared to both the true and predicted regions in simulations with different migration times.

First, we wanted to see whether there is a difference in the length distribution of false positive regions compared to true ones.

We hoped that the lengths of predicted introgressed Hum→Nea elements might be informative for the time of migration. However, there is more power to find longer elements, and this ascertainment bias is so strong that information about the timing of migration is almost completely lost. Overall, our predicted elements were somewhat longer

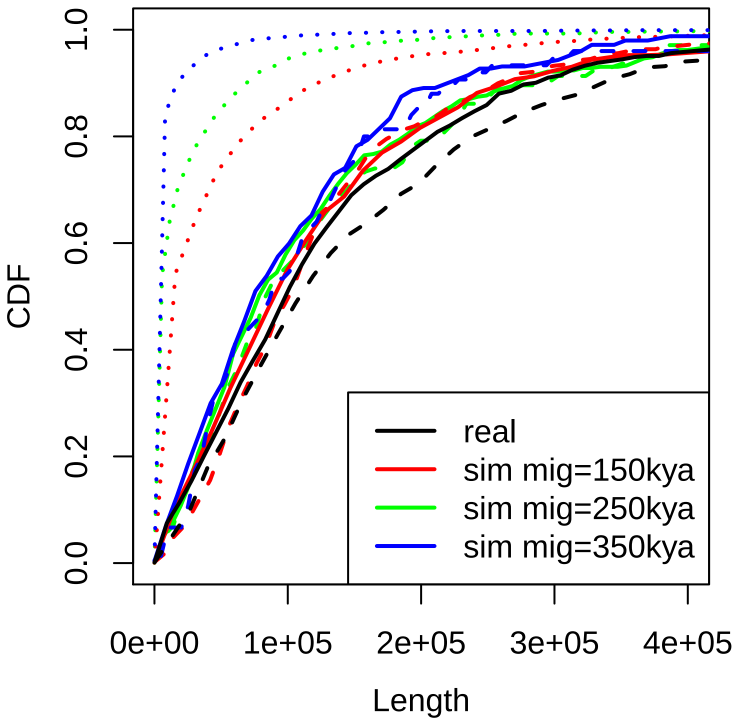


Figure S6: The distribution of lengths in real vs simulated Hum→Nea regions. The dotted lines show the true distribution of lengths in three simulated data sets, each produced with different migration times, indicated by the color. The solid lines show the distribution of regions found by ARGweaver-D in the simulated and real (black) data sets when analyzed with a model with a migration band at 250kya. The dashed lines show the same except with a migration band at 150kya.

(median 105kb) than those observed in our simulations (median lengths ranging from 71kb-89kb), but the distributions were largely overlapping (Supp Fig S6). While all the ARGweaver-D analysis was done with a simple constant recombination model, the underlying recombination map is also an important factor, and little is known about the Neanderthal recombination map.

#### Validation of super-archaic regions in SGDP individuals

We further explored the 27Mb of the genome which was putatively identified as Sup→Den. This category had the strongest prior evidence for super-archaic introgression [5, 6], and is the only super-archaic category for which the amount detected by ARGweaver-D (1%) significantly exceeds the false positive rate estimated from simulations (0.5%). We first identified variants in Sup→Den regions that map to the migrant lineage in our data set (which included the Denisovan, two Neanderthals, SGDP individuals Khomani_San_1 and Mandenka_2, and chimpanzee); there are 15,470 variants over 16.8Mb of unmasked Denisovan sequence. This suggests an average substitution rate is 9.2e-4/bp, which translates to a branch length of 1.8My (using a mutation rate of 1.45e-8/bp/generation and a generation time of 29 years). However, we expect that this estimate is biased upwards, as Sup→Den regions with more variants are easier to detect. We compiled a VCF file containing all the substitutions on the super-archaic haplotype.

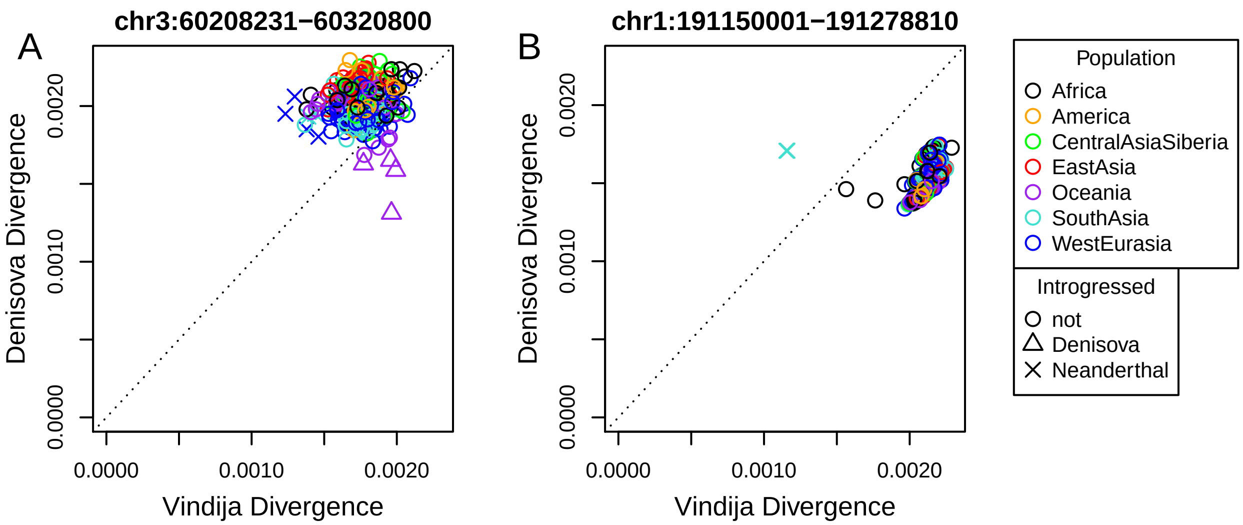


Figure S7: **Average divergence of SGDP individuals to Neanderthal and Denisovans in example Sup→Den regions** The color represents the population of each sample, whereas the symbol indicates whether introgression has been detected in this individual in at least half of the plotted region (according to the CRF method). A) This region shows the expected pattern for Sup→Den: most individuals have higher divergence to the Denisovan than to the Vindija Neanderthal. However, a few Oceanian individuals who have Denisovan introgression in this region have lower Denisovan divergence. Similarly, some European individuals with Neanderthal introgression also show a decreased Neanderthal divergence. B) This is a less typical Sup→Den region that is likely a false positive, as most SGDP individuals show lower divergence to the Denisovan than to Neanderthal. It is interesting that one of the outlying African dots represents an individual used in the ARGweaver-D analysis (Khomani_San_1).

We next looked at all 279 individuals in the SGDP data set, comparing their divergence to Neanderthals and Denisovans in each region. If the Sup→Den prediction is correct, then the Denisovan divergence should be high for all humans, and not just the two humans used in the ARGweaver-D analysis. Two example regions are shown in Figure S7. As described in the caption, and explored further below, most Sup→Den regions do indeed show higher divergence to the Denisovan across the SGDP individuals than to the Neanderthal, excepting individuals with Den→Hum introgression in that region. There are a small number of Sup→Den regions, such as the one shown in Figure S7B, where the Denisovan divergence does not exceed Vindija divergence in most humans and might be false positives of the ARGweaver-D approach.

While looking at example plots is helpful, we want to summarize the properties across all Sup→Den regions. We define a statistic $f$, which is the fraction of SGDP individuals in a given region for which the Denisovan divergence is greater than the divergence to either Neanderthal. This statistic can be visualized as the fraction of individuals that fall above the diagonal in plots such as those in Figure S7. In each region we exclude any individuals with Neanderthal or Denisovan introgression (as assigned by the CRF [7]). We computed $f$ for each of the 161 putative Sup→Den regions with length $\geq$ 50kb, as well as for 262 putatitve Sup→Nea regions, 384 putatitve Sup→Afr regions, and 500 100kb regions randomly selected from regions of the genome without any ARGweaver-D introgression assignment. The distributions of $f$ for each set of regions are shown in Figure S8. We see that for about 80% of the Sup→Den

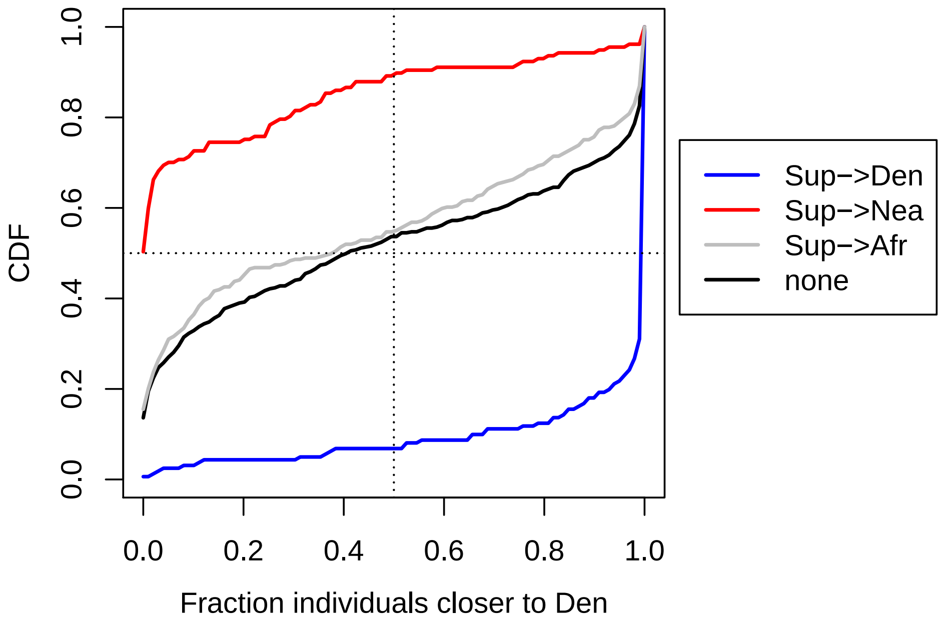


Figure S8: **Distribution of the fraction of individuals with higher Denisovan vs Neanderthal divergence**. Each cumulative distribution shown here is taken across a set of regions assigned to a particular introgression category by ARGweaver-D. For each region, we calculate the number of individuals for which the Denisovan divergence is higher than the Neanderthal divergence (excluding individuals with calls of Neanderthal or Denisovan introgression by the CRF). We see that Sup→Den regions have a high proportion of individuals more closely related to Neanderthal, and the opposite pattern in Sup→Nea regions. Both putative Sup→Afr and non-introgressed regions are very slightly biased towards Neanderthal ancestry.

regions, $f$ is close to 1. There are 29 Sup→Den regions with $f < 0.9$ (including the region shown in Figure S7B), and which might be best regarded as false positives.

For the Sup→Nea regions, where we would expect most individuals to have higher Neanderthal than Denisovan divergence, we see a similar large shift towards small $f$; most SGDP individuals are closer to Denisovans than Neanderthals in these regions. In this case, 73% of the regions have at least 90% of individuals closer to the Denisovan.

It is important to note that while this analysis provides a check on ARGweaver-D's predictions, and identifies some potential false positives, it does not imply that the remaining regions are true positives. Other scenarios, such as balancing selection, could also produce long regions of high divergence that may be virtually indistinguishable from super-archaic introgression. But this analysis does show that the signal identified using only two humans usually holds across a much larger set.

#### Analysis of Sup→Den regions passed to modern humans

Presumably, if there is super-archaic introgression into Denisovans, and later Denisovan introgression into Oceanian and Asian humans, then it seems likely that these modern humans harbor super-archaic alleles passed through the

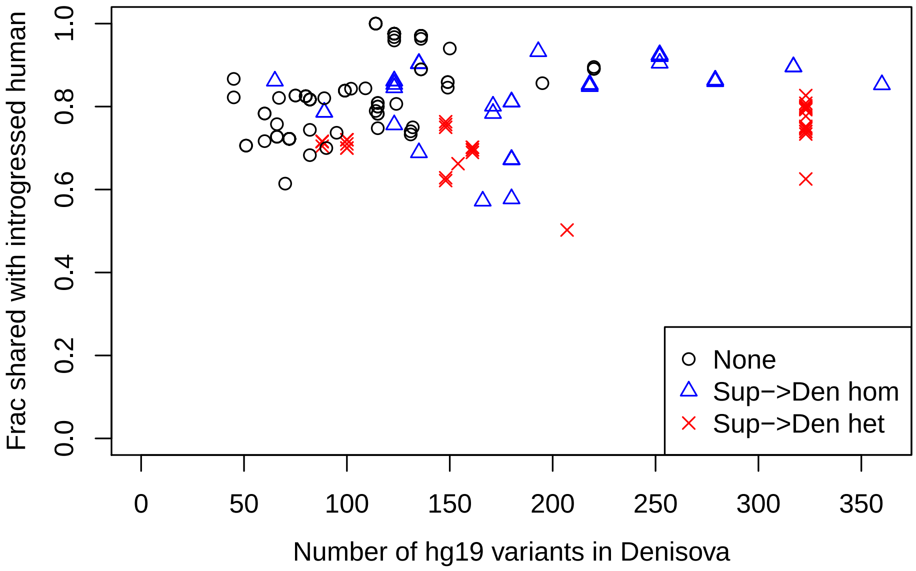


Figure S9: **Fraction of shared variants with Denisovan, for individuals with Denisovan introgression.** Each point is calculated for a particular genomic region and individual with Denisovan introgression in that region. The x-axis shows the number of Denisovan hg19 differences; the y-axis shows the fraction of these variants shared with the individual. The colors represent the type of region; blue regions are homozygous Sup→Den regions, red regions are heterozygous Sup→Den regions, and black are regions without any ARGweaver-D introgression calls in Africans or archaics.

Denisovans. Indeed, 15% of our Sup→Den regions overlap regions with Den→Hum introgression calls in the SGDP (24 out of 161 regions, excluding regions with lengths <50kb). We looked into this by comparing the variants on the super-archaic lineage with those observed in individuals with Denisovan introgression (according to the CRF calls). Figure S9 shows the fraction of shared Denisovan variants vs. the number of hg19/Denisovan variants for individuals that are annotated with Hum→Den introgression by the CRF method.

The black points in Figure S9 show the fraction of shared variants in regions without any ARGweaver-D introgression calls in Africans or archaics. We see that the fraction of shared alleles is high (between 60-100%) for these regions, though the overall number of variants is moderate. The blue points show the same values in regions that have been identified as Sup→Den introgressed in both Denisovan lineages. For the most part, we also see high fraction of shared variants, although the absolute number of variants is much higher overall. This suggests that the individual is sharing super-archaic alleles, as the majority of these variants occur on the super-archaic branch. Finally, the red points show a subset of Sup→Den where the super-archaic introgression is only found in one of the Denisovan lineages, so that our sampled Denisovan has both a ”super-archaic” and a ”Denisovan” haplotype. The red points with the lowest fraction shared may represent individuals who received the Denisovan haplotype.

One consideration here is that there is likely a bias towards identifying Denisovan introgression in humans when the Denisovan and human both share the super-archaic haplotype, because introgression will be very easy to detect with such a large number of shared variants. This may explain why there are not many red points with lower fractions of shared Denisovan variants in Figure S9. Regardless, this analysis shows that many individuals with Denisovan introgression share alleles that are predicted introgressed into Denisovan from a super-archaic hominin.

#### Analysis of Sup→Nea regions passed to modern humans (and the hg19 reference sequence)

We did a similar analysis on regions identfied as Sup→Nea, this time looking at the overlap between these regions and Nea→Hum regions in SGDP humans. 35% of our Sup→Nea regions overlap regions with Nea→Hum introgression according to the CRF predictions (55 out of 157 regions, excluding regions with length <50kb). Figure S10 summarizes these regions; there are many more points in this plot than in Figure S9 because there are many more SGDP samples with Neanderthal introgression.

The first surprising aspect of these results is that there was one region (chr6:8450001-8563749) classified by ARGweaver-D as Sup→Nea, but which had only 13 hg19 differences across 79kb of unmasked Neanderthal sequence (giving an hg19/Neanderthal divergence of only 0.016%). After closer inspection, we suspect that this region has Neanderthal introgression in the hg19 reference sequence. Among SGDP individuals with annotated Neanderthal introgression in this region, there are between 1 and 18 homozygous hg19 differences. Among the other SGDP individuals, there are between 68 and 369 homozygous hg19 differences. (This excepts one individual, Saharawi_1, fosr whom CRF introgression calls were not made and which has only one hg19 difference; we presume it is also Neanderthal introgressed. The Saharawi are a Northern African population that has been found to have almost as much Neanderthal introgression as non-African populations [6]). Given that the reference sequence is largely African-American, we should expect that it would contain some Neanderthal ancestry. While the theory of possible introgression from super-archaic introgression into Neanderthal does not yet have strong support, if the annotated Sup→Nea regions are correct, this would be an example of archaic hominin ancestry in the hg19 reference sequence, passed through Neanderthals. We also note that we found one other region (chr10:88106371-88206370), from our set of 500 randomly selected 100kb regions without ARGweaver-D introgression calls, in which the number of hg19 differences is $< 4$ for SGDP individuals with Neanderthal introgression, and much higher (median: 75) for other SGDP individuals. As our analysis in this section only spanned 2.4% of the genome, a genome scan for Neanderthal introgression on hg19 would discover many more such regions.

Beyond the observation of Neanderthal (and possibly super-archaic) ancestry in hg19, Figure S10 shows that there are indeed a number of regions annotated as Sup→Nea with a large number of hg19 variants, which are also shared to a high degree with Neanderthal-introgressed humans in the SGDP. Again, we see that the red points (representing non-fixed Sup→Nea regions) sometimes share fewer variants with Neanderthal, suggesting that these individuals are introgressed with the 'Neanderthal' haplotype rather than the 'super-archaic' haplotype.

Overall, the analysis of both Sup→Den and Sup→Nea regions show that these regions have a high number of variants compared to non-annotated regions, and that these variants are often shared with humans with introgression from Denisovans or Neanderthals (respectively). While the Sup→Nea event is not well supported, and the Sup→Nea regions may simply be highly diverged Neanderthal regions, there is stronger support for the Sup→Den migration, and

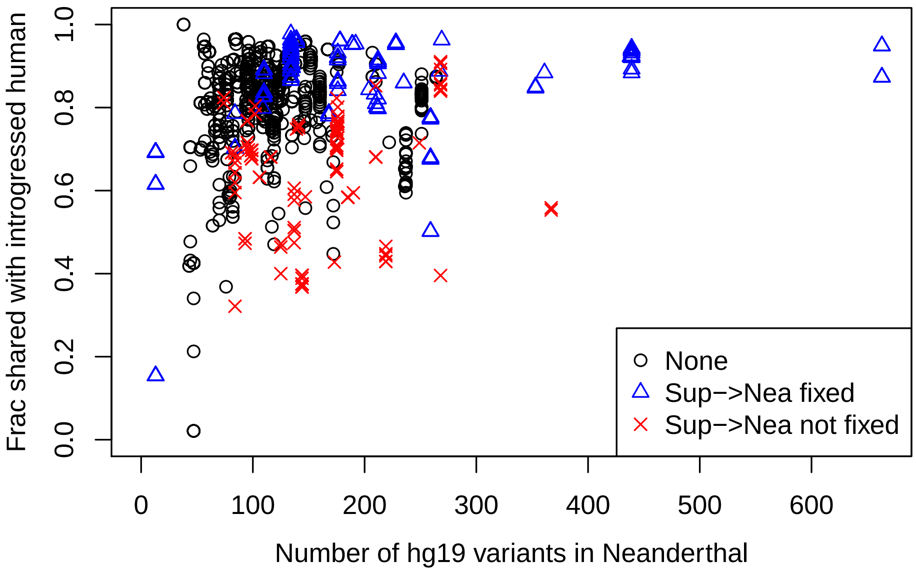


Figure S10: **Fraction of shared variants with Neanderthal, for individuals with Neanderthal introgression.** Each point is calculated for a particular genomic region and individual with Neanderthal introgression in that region. The x-axis shows the number of Neanderthal hg19 differences in the region; the y-axis shows the fraction of these variants shared with the individual. The colors represent the type of region; blue regions are Sup→Nea regions, red regions are a subset of Sup→Nea that are not fixed in our Neanderthal sample, and black are regions without any ARGweaver-D introgression calls in Africans or archaics.

it seems that humans with Denisovan ancestry must also harbor some variants from more diverged hominin species as well.

#### Functional enrichment analysis of introgressed regions

We checked for enrichments or depletions of various functional elements in our introgressed segments. However, the interpretation of these numbers is not straightforward, as the power to detect the segments is confounded by many factors, such as local population size, mutation rate, recombination rate, and sequence quality filters. Indeed, even for introgression into humans where the detection power is quite high, these depletions have been difficult to interpret (see Discussion). The enrichments are shown in Supp Fig S11. It is clear that biases for functional elements depend on the type of introgression event (from a sample population or super-archaic). We found a 1.15x enrichment of ensembl CDS regions in our Hum→Nea calls, which is most likely explained by higher power with lower effective population size. Perhaps the most interesting aspect is that the enrichment Hum→Nea in most functional categories (including CDS, phastCons, promoters) is larger in the Vindija Neanderthal than the Altai, despite the fact that the Vindija and Altai Hum→Nea regions are highly overlapping (55.6% of the combined set are called in both individuals). Again, it appears that functional elements were not lost in the duration between the Altai and Vindija individual's lifetimes, suggesting an absense of negative selection acting against these regions.

#### Deep introgression analysis with other models

In the main paper, we presented results using an ARGweaver-D model in which all migration times ($t_{mig}$) were set to 250kya and super-archaic divergence time ($t_{div}$) to 1Mya. We also ran ARGweaver-D genome-wide with $t_{mig}$ =150kya and $t_{div}$=1.5kya. The coverage of the resulting elements is shown in Figure S12.

The results using this model are qualitatively similar to those presented in the main paper; the largest coverage is in Hum→Nea, with increased coverage on the X chromosome, and a somewhat smaller depletion from Altai to Vindija on the X. We again see similar low levels of all other introgression categories, and the same depletion for Sup→Den on the X chromosome. The coverages of predicted introgressed Hum→Nea and Sup→Den elements are about half what is presented in the main paper. This is consistent with our simulation results which show that power is much lower using this model when the true model more closely matches the one used in the paper, and further supports our claims that the Hum→Nea migration was quite old, and that the super-archaic divergence was low.

We also did a genome-wide run in which we included the Papuan and Basque, along with the two Africans and archaic individuals. We analyzed the data as before, using $t_{mig}$ = 250kya and $t_{div}$ = 1Mya, but also added migration bands from Nea→Hum and Den→Hum at 50kya, for a total of 7 migration bands. The introgression coverages for this run are presented in Supplementary Figure S13. Although there are moderate differences in the absolute level of introgression predicted, the results agree well with those presented in the main paper. We do see an increase in the amount of Hum→Den and Hum→Nea regions, which most likely can be explained by the algorithm calling the incorrect direction of migration for some instances of Den→Hum and Nea→Hum.

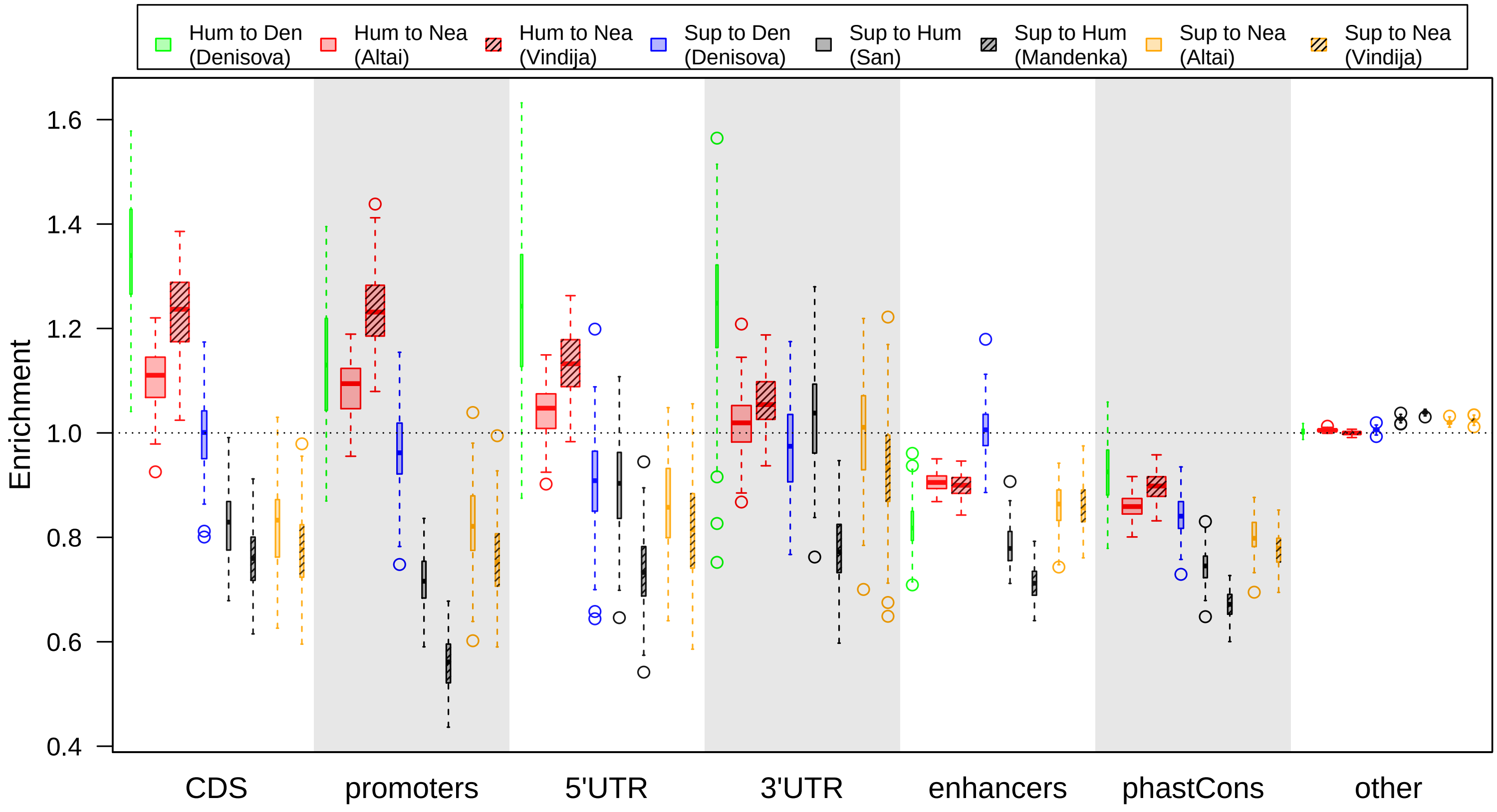


Figure S11: **Enrichment of predicted introgressed regions within different annotation groups.** Enrichment is calculated as the number of introgressed bases called within a particular category, divided by the number expected assuming that the introgression calls are independent of the annotations. The distributions shown in the box plots are calculated from 100 bootstrap replicates over the introgression calls. The width of each bar is proportional to the total coverage of the predicted introgressed elements genome-wide. These enrichments are likely highly influenced by factors affecting the power and false positive rates of the calling algorithm (See discussion in Supplementary Section XX).

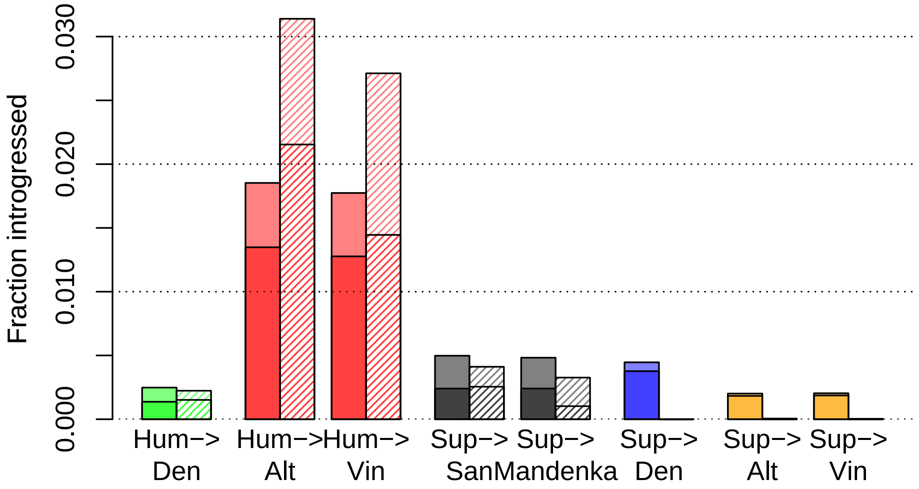


Figure S12: **Introgression coverages under alternative demographic model**. Genome-wide coverages using a migration time of 150kya and a super-archaic divergence of 1.5Mya. Solid bars show autosomal coverage; striped bars show X-chromosome coverage. The dark bottom halves of each bar represent regions that were also predicted by the model used in the main paper.

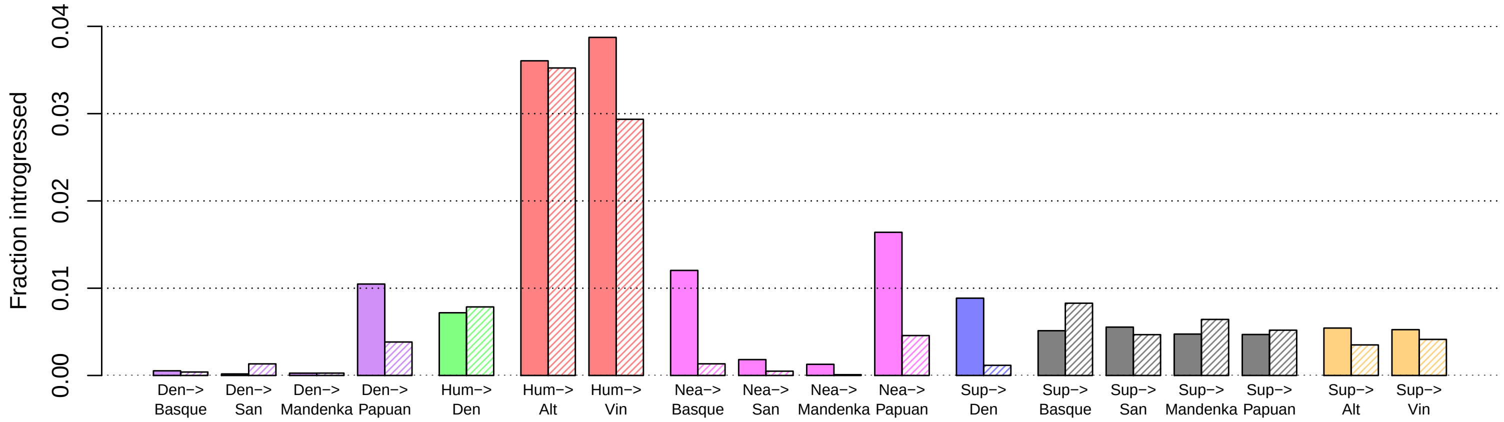


Figure S13: **Supplementary Figure: Coverages using out-of-Africa model and full population tree. Notation as in Figure S12, with the addition of Papuan and Basque individuals.**

##### Fraction of Neanderthal genome introgressed from humans

Using the true and false positive rates in from the deep introgression simulations in the main paper, we can make a rough estimate of the total amount of Neanderthal genome introgressed from ancient humans. If the fraction of the genome predicted to be introgressed is $x$ , then the total amount is predicted to be $(x - FP)/TP$. These numbers are of course very rough because there are many unknown demographic parameters in the model. Using the results in the main paper, this would predict that 7.4% of the Altai autosomal genome is introgressed from Neanderthal, and 7.2% of the Vindija. If we instead use the simulation and real results from the previous supplemental section, where the demographic model has a migration at 150kya, we get predictions of 10.8% for Altai and 10.3% for Vindija.

#### Calculating the mutation rate map

We first extracted the hg19 (human), panTro4 (chimp), gorGor3 (gorilla), ponAbe2 (orangutan), and nomLeu3 (gibbon) sequences from the UCSC Genome Browser's 100-way vertebrate alignment. We then masked any segments of the alignment within 100bp of a phastCons element, using the union of all hg19 phastCons elements (phastConsElements46way, phastConsElements46wayPlacental, phastConsElements46wayPrimates, phastConsElements100way). We then ran phyloP on the alignment, using the options `--method LRT --features windows.bed --mode CONACC`, where the windows.bed file gives 100kb sliding windows of the human genome, staggered by 10kb. The substitution model used for the phyloP run was downloaded from `http://hgdownload.soe.ucsc.edu/goldenPath/hg19/phastCons100way/hg19.100way.phastCons.mod`. The phyloP run produced tree scaling factors for every 100kb window. These were converted to mutation rates by rescaling all factors to achieve a mean mutation rate of 1.45e-8/bp/generation. The mutation rate for a particular base was then mean of the mutation rates in the sliding windows which overlap that base. For substitution rates on chromosome X, we used the same procedure, with the exception that we used a substitution model specific to chromosome X, downloaded from `http://hgdownload.soe.ucsc.edu/goldenPath/hg19/phastCons46way/primates.chrX.mod`.
